# Hidden grassland productivity from overlooked herbivore impacts

**DOI:** 10.64898/2026.10.13.758807

**Authors:** Yan Shi, Hao Wang, Hongbiao Zi, Xiangsheng Wang, Huijie Gu, Zhenhua Zhang, Liqing Peng, Anping Chen, Guirui Yu, Jin-Sheng He

**Author notes:** These authors contributed equally: Yan Shi, Hao Wang.

## Abstract

Grasslands cover approximately 40% of Earth’s land surface and contribute substantially to terrestrial primary production. Despite widespread grazing, field estimates of grassland aboveground net primary productivity commonly overlook two grazing-related processes: biomass consumed by herbivores and post-grazing compensatory growth. The magnitude and direction of the resulting biases remain unclear. Here we quantified these biases using a 3-year yak grazing experiment on the Tibetan Plateau, a meta-analysis of 152 grazing experiments across China, and a global synthesis of 212 paired observations from 58 studies. Overlooking biomass consumed by herbivores consistently underestimated productivity by 34–51%. By contrast, overlooking grazing-induced compensation generated smaller but more variable biases, ranging from 29% underestimation to 19% overestimation, mainly reflecting contrasting compensatory responses under rotational versus continuous grazing. These findings show that overlooking grazing-mediated processes introduces substantial, process-dependent biases in grassland productivity estimates, underscoring the need to incorporate these processes into ecosystem and climate assessments.

## Introduction

Grasslands cover about 40% of Earth’s land surface and account for 69% of global agricultural land use, making them one of the most extensive and ecologically important biomes on the planet^1^. Through long-term plant–herbivore coevolution, large herbivores strongly shape grassland ecosystems by grazing, trampling, and redistributing nutrients, thereby affecting biodiversity, carbon cycling, and ecosystem multifunctionality^2,3^. One important effect is their direct influence on aboveground net primary productivity (ANPP), a core process underpinning carbon inputs, energy flow, forage supply, and ecosystem functioning in grazed landscapes^4,5^.

Yet ANPP in grazed grasslands is commonly estimated in ways that omit herbivory itself. Field methods typically either measure peak biomass in grazed plots, thereby missing biomass already removed by animals, or infer productivity from ungrazed exclosures, thereby missing grazing-induced regrowth and, in some cases, overcompensation^6–8^. We refer to these two sources of bias as overlooked animal consumption (OAC) and overlooked growth compensation (OGC) (Fig. 1a and Supplementary Fig. 1). This problem is conceptually important because ANPP is a production flux, whereas peak standing biomass is a state variable^9,10^. In grazed systems, herbivory decouples the two by both removing produced biomass and altering subsequent regrowth, so standing biomass no longer provides a reliable proxy for annual biomass production^8,11^. The magnitude and even direction of this bias are likely to vary because compensatory responses depend on grazing intensity, seasonality, and management regime^6,12,13^, ranging from overcompensation under moderate, rotational grazing to undercompensation under chronic heavy grazing^14–17^. Remote sensing does not resolve this problem either: vegetation indices such as Normalized Difference Vegetation Index (NDVI) largely capture standing biomass, not the production hidden in consumed tissue or post-grazing recovery^18,19^. More broadly, recent syntheses of ecosystem carbon cycling suggest that animals remain underrepresented in carbon accounting^20^, even though excluding herbivory can alter estimates of ecosystem carbon uptake and storage.

**Fig. 1.**
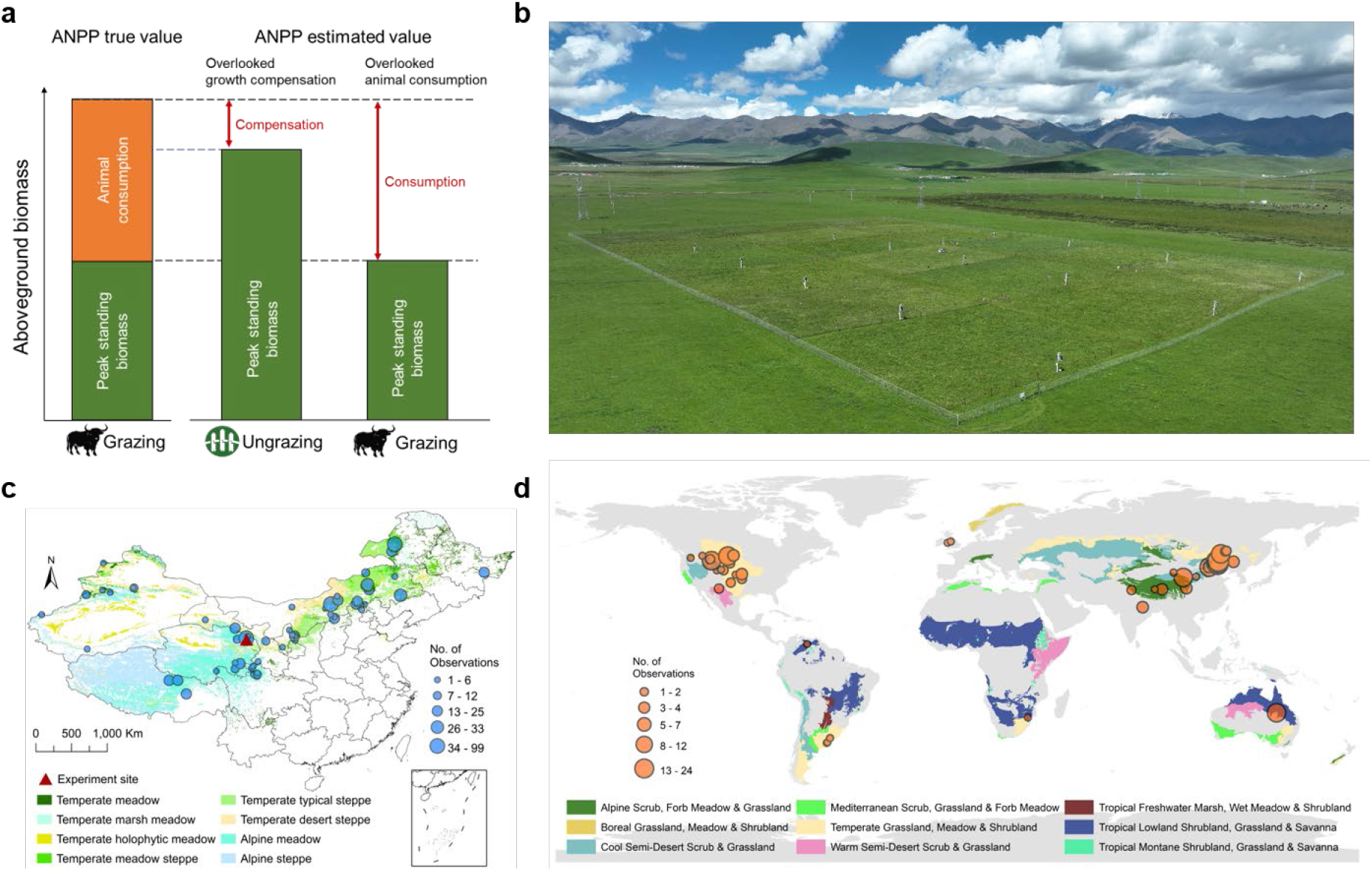
Grazing grassland ANPP estimation error: conceptual framework, the validated experiment, and synthesis analysis. **a**, Conceptual framework illustrating ANPP estimation error from traditional peak standing biomass measurements in grazing and ungrazing grasslands. **b**, Aerial view of the grazing experiment on the Tibetan Plateau. **c**, Locations of grazing experiments (152 studies) in China included in the meta-analysis. **d**, Locations of grazing experiments (58 studies) for global synthesis.

Here we quantify these biases across scales using a 3-year yak grazing experiment in Tibetan alpine grasslands (Fig. 1b), a meta-analysis of 152 grazing experiments across China (Fig. 1c), and a global synthesis combining five published ANPP datasets for OAC and 212 field observations from 58 studies for OGC (Fig. 1d). We show that herbivory significantly alters the accuracy of field-based grassland productivity estimates, beyond its effects on productivity itself. By separating production lost to consumption from production regained through compensatory growth, we identify a pervasive source of bias in grazed-grassland ANPP estimates and provide a basis for improving ecological inference, carbon accounting, and grassland monitoring worldwide.

## Results

### Experimental evidence

The yak grazing experiment shows that conventional ANPP estimates can miss a substantial fraction of production. Across the 3-year experiment (2021–2023) (Supplementary Figs. 2 and 3), estimating ANPP from peak biomass in grazed plots underestimated alpine grassland productivity by ∼51% on average because biomass consumed by yaks was excluded from the estimate (OAC method; Fig. 2; Supplementary Tables 1–3). Estimating ANPP from ungrazed controls also underestimated productivity, by ∼25% on average, because grazing-induced compensatory response was ignored (OGC method). The two standard field approaches therefore failed for opposite biological reasons: one overlooked biomass already removed by herbivores, whereas the other treated ungrazed growth as an unbiased reference for grazed plants.

**Fig. 2.**
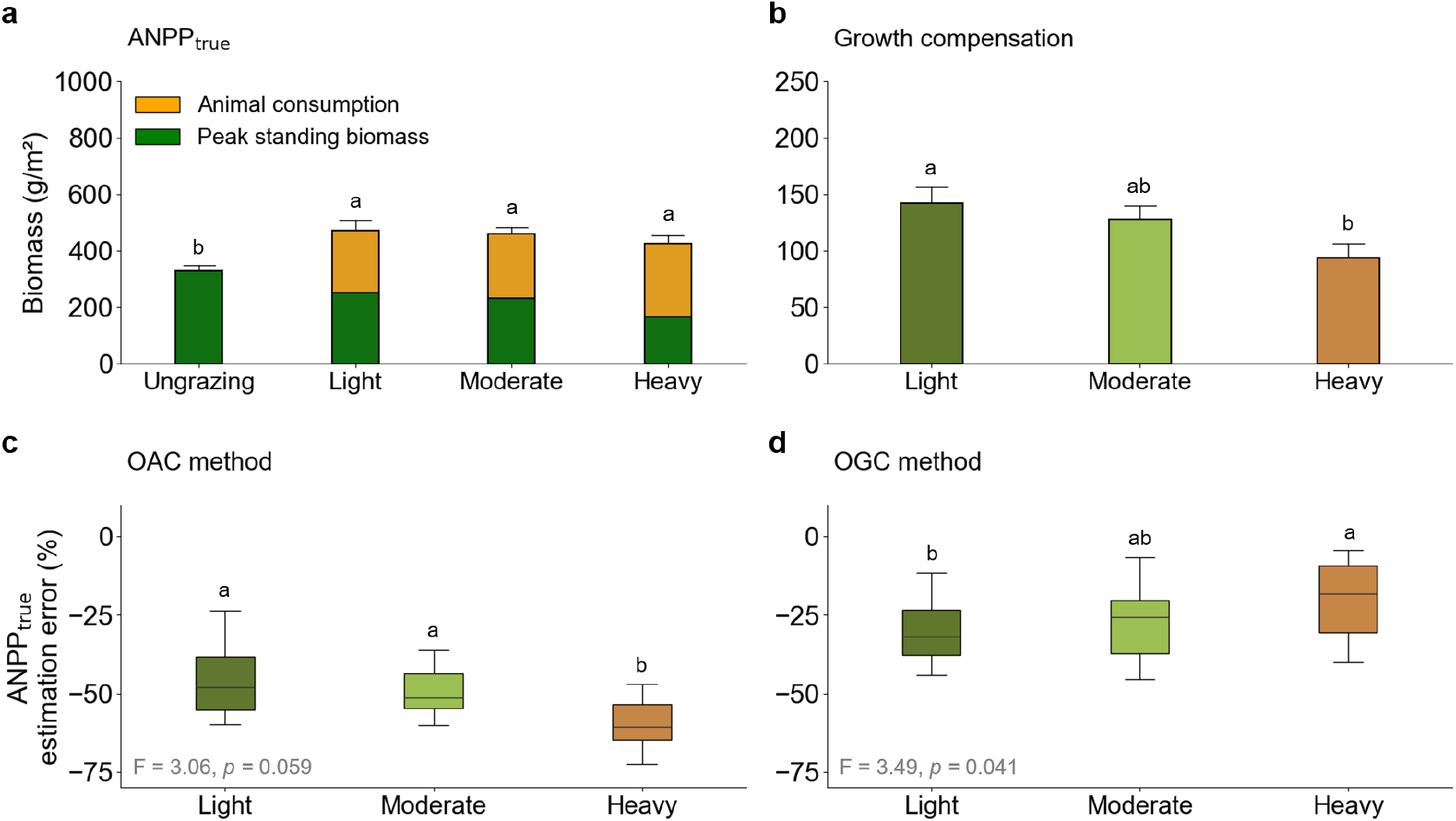
Effects of grazing on ANPP and its estimation error in the grazing experiment. A and **b**, ANPP_true_ components under different grazing intensities: standing and consumed biomass (**a**) and growth compensation (**b**). **c** and **d**, Estimation errors of ANPP_true_ in grazing plots by the OAC (Overlooked animal consumption) method (**c**) and the OGC (Overlooked growth compensation) method (**d**), respectively. Positive values signify overestimation and negative values denote underestimation. Different letters indicate significant differences among treatments at *p* < 0.05.

**Fig. 3.**
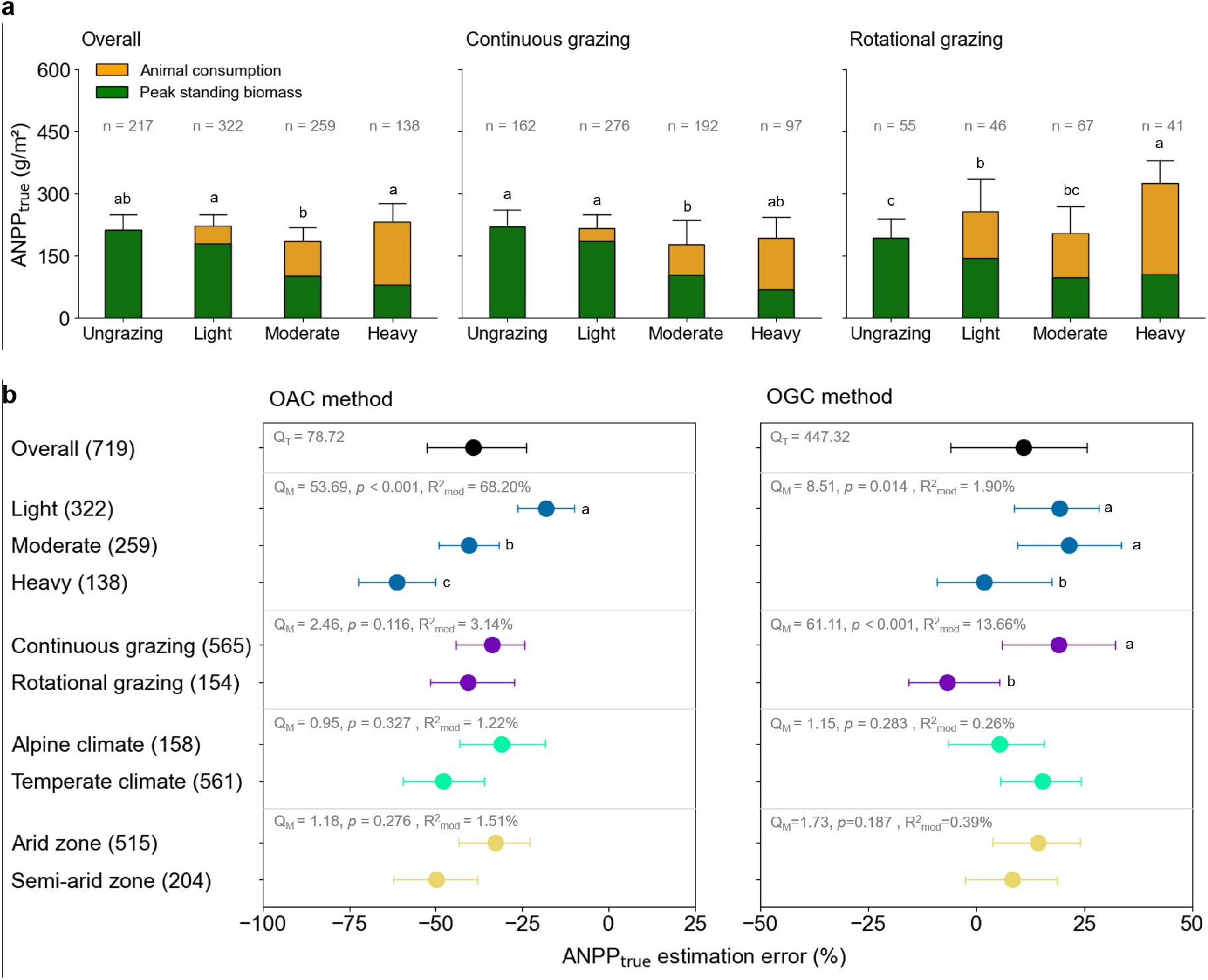
Effects of grazing on ANPP and its estimation error from a meta-analysis of northern China grasslands. **a**, ANPP components and their responses to grazing intensities and regimes. **b**, ANPP estimation error by OAC (Overlooked animal consumption) and OGC (Overlooked growth compensation) method across all studies and by categories of grazing intensity, grazing regime, and climate type. Locations of grazing experiments included are illustrated in Fig. 1c. Positive values indicate overestimation and negative values indicate underestimation. Numbers in parentheses denote sample sizes.

The two biases responded differently to grazing intensity. OAC-related underestimation increased from ∼45% under light grazing to ∼58% under heavy grazing, tracking greater biomass removal as stocking pressure rose (Fig. 2a, c). By contrast, OGC-related underestimation declined from ∼29% to ∼21% with increasing grazing intensity, as growth compensation weakened (Fig. 2b, d). Similar magnitudes have been reported in other alpine meadow experiments on the Qinghai-Tibetan Plateau, where medium and high yak stocking consumed standing herbage biomass by about 15% and 27% relative to ungrazed controls, while relative compensation under moderate grazing typically ranged from 20 to 60%^21–24^. Together, these results show that herbivory introduces no single correction to ANPP estimates: consumption bias strengthens as grazing intensifies, whereas compensatory gains weaken.

This contrast is consistent with the ecology of alpine grasslands. Under light to moderate grazing, defoliation can still be followed by partial regrowth, but as grazing pressure increases, repeated tissue removal increasingly outpaces recovery in a system already constrained by a short growing season and limited recovery time^15,24^. Peak standing biomass in grazed alpine meadow therefore becomes a progressively poorer proxy for ANPP, not due to random measurement noise, but because grazing alters both how much biomass is removed and how much production is regained after defoliation.

### Patterns across Chinese grasslands

The broader Chinese synthesis shows that the experimental pattern is reproducible across a much wider range of environmental and management gradients (Fig. 3 and Supplementary Table 4). Across 936 observations from 152 studies spanning eight grassland types, OAC caused an average underestimation of ANPP of about 38% (Fig. 3b). Most of the heterogeneity in this bias was explained by grazing intensity, with biomass consumed by animal and underestimation increasing as intensity increased (Fig. 3a, b). This national-scale pattern closely mirrors the yak experiment and indicates that biomass removal is the more regionally consistent source of ANPP bias in grazed grasslands. More generally, studies across China show that aboveground biomass and productivity often remain relatively unchanged under light grazing but decline progressively as stocking pressure increases, particularly where a larger fraction of current-year growth is removed before peak biomass is sampled^21^.

OGC showed a different structure. Across the Chinese dataset, ignoring compensatory growth produced an average overestimation of ANPP of about 12% (Fig. 3b and Supplementary Table 4). Here, both grazing intensity and grazing regime mattered. Light and moderate grazing generated larger overestimation responses than heavy grazing, consistent with the grazing optimization hypothesis, which predicts a unimodal relationship between grazing intensity and productivity, with maximum productivity occurring at moderate grazing^25,26^. Grazing regime also changed the direction of bias. Under rotational grazing, ANPP was underestimated by about 7% relative to ungrazed plots, consistent with overcompensatory regrowth, whereas under continuous grazing, ANPP was overestimated by about 19%, consistent with undercompensation (Fig. 3a, b). This contrast likely reflects constrained post-grazing recovery under continuous grazing, which reduces the capacity of vegetation to replace consumed tissue^27^.

Consistent with our yak grazing experiment on the Tibetan Plateau, rotational grazing can maintain or even increase aboveground biomass relative to ungrazed plots, indicating that compensatory growth can partially offset tissue loss under favorable grazing regimes ^28^. Yet plateau-wide synthesis also shows that this recovery weakens under heavier grazing, when declines in biomass, soil resources, and plant recovery capacity become more pronounced ^26^. Across Chinese grasslands, grazing intensity therefore primarily controls consumption bias, whereas grazing regime and intensity jointly regulate regrowth bias. OAC tracks how much production is removed; OGC reflects not only biomass removal but also whether local management permits vegetation recovery after defoliation. This distinction helps explain why OAC is easier to generalize across regions, while OGC remains more contingent on local ecological conditions.

### Global bias in grassland ANPP estimates

At the global scale, overlooked animal consumption remained a major source of error. By integrating published datasets on peak standing biomass, livestock populations, foraging duration, and grassland distribution, we estimate that OAC causes a mean underestimation of global grassland ANPP of about 34% (Fig. 4a; Supplementary Table 5). The magnitude of this bias varied strongly among regions and grassland types, reaching about 49% in savannas and about 44% in North American grasslands, but only about 15% in alpine grasslands (Fig. 4b–f). ANPP underestimation also increased from cold to warm and from arid to humid climates (Fig. 4g, h), indicating that the largest hidden productivity occurs in warm, humid grasslands, where high primary production supports greater herbivore consumption^29,30^. This climatic pattern is consistent with broader views of grassy biomes as systems shaped by interacting controls of climate, soils, fire, and herbivory rather than by climate alone^3^.

**Fig. 4.**
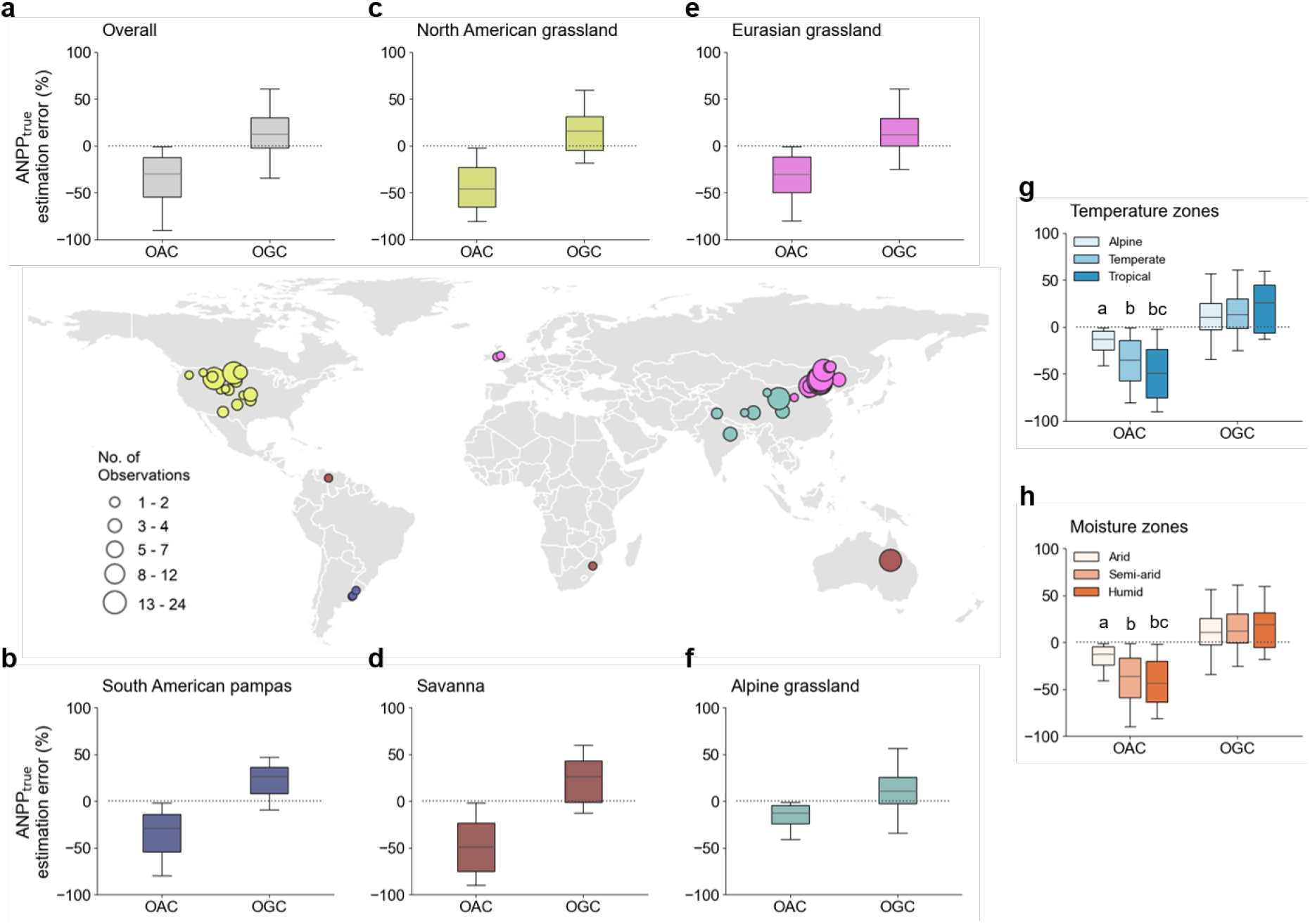
Global synthesis of ANPP estimation errors by OAC and OGC in grazed grasslands. **a**, Overall ANPP estimation error. **b–f**, ANPP estimation error for South American pampas (**b**), North American Prairie (**c**), savanna (**d**), Eurasian grassland (**e**), and alpine grassland (**f**). **g** and **h**, ANPP estimation error in different temperature (**g**) and moisture zones (**h**). Different letters denote significant differences among climate zones at *p* < 0.05. Positive values indicate overestimation and negative values indicate underestimation.

These spatial differences suggest that global OAC is structured primarily by the coupling between productivity and biomass removal. In warm, humid, and highly productive grasslands, larger amounts of biomass can be produced and consumed before peak standing crop is measured, widening the gap between observed biomass and true ANPP. By contrast, in alpine grasslands, shorter growing seasons and lower productivity constrain both removal and recovery, limiting the amount of hidden production. The largest OAC biases should therefore be expected not in the most environmentally constrained systems, but in the most productive grazed systems, where herbivores can remove a substantial fraction of seasonal growth before sampling. In this sense, OAC is not simply a livestock effect; it is a broad-scale productivity-dependent bias in how ANPP is inferred from standing biomass.

Global OGC showed a different structure. Using 212 observations from 58 studies under continuous moderate grazing, we estimated a mean ANPP overestimation of about 14% when grazing-induced compensatory responses were ignored (Fig. 4a; Supplementary Table 5). Unlike OAC, this bias did not vary significantly among ecozones or broad climate classes (Fig. 4g, h), suggesting that compensation bias is governed less by broad climatic gradients than by localgrazing regime, grazing history, and plant recovery capacity ^15,16,31^. This asymmetry indicates that OAC is the more spatially predictable component of herbivory-induced bias, because it mainly reflects biomass removal, whereas OGC additionally depends on whether grazing still permits compensatory growth. As a result, global OAC-related ANPP underestimation is likely to be strongest in warm, productive grasslands with high herbivore pressure, while OGC-related error may remain more heterogeneous and management-dependent even within similar climate zones^32–35^.

The contrast between OAC and OGC also has implications for large-scale prediction. OAC can be approximated from broad-scale information on biomass, livestock abundance, and grazing duration because it tracks how much production is removed. OGC is harder to generalize because compensatory growth depends on local recovery conditions that are not captured by coarse climate classes alone. The weak climatic structure of OGC therefore does not imply that regrowth is unimportant; instead, it shows that broad-scale global patterns are better suited to predicting consumption bias than recovery bias. This makes OAC the more tractable target for first-order correction of global ANPP estimates, whereas improving OGC estimates will require finer information on grazing management, vegetation state, and recovery potential.

## Discussion

These results show that herbivory is not only an ecological driver of grassland function but also a source of systematic measurement bias. Accurate ANPP estimates are central to understanding ecosystem services, carbon turnover, and rangeland management^7^. By combining experimental, national, and global evidence, we show that conventional ANPP estimates in grazed grasslands can be biased downward when animal consumption is ignored and biased either downward or upward when compensatory growth is ignored. Herbivory therefore affects both the process being measured and the validity of the measurement itself.

The two biases differ in both generality and consequence. OAC is the more consistent signal: wherever grazers remove a substantial fraction of seasonal growth, peak standing biomass will underestimate production. This helps explain why true ANPP in grazed systems can greatly exceed standing biomass and why underestimation strengthens with grazing intensity in both our experiment and broader syntheses^15,24^. OGC is less spatially predictable but no less important. Because compensatory growth depends on grazing regime, grazing history, and plantrecovery capacity, ungrazed exclosures do not provide a neutral benchmark in working grasslands ^13,36,37^. The same field protocol can therefore underestimate ANPP where grazing stimulates regrowth and overestimate it where repeated defoliation suppresses recovery.

These biases matter because ANPP is widely used well beyond productivity studies. If consumed biomass is omitted, forage supply and carrying capacity will be underestimated; if compensatory responses are misrepresented, the productivity of managed grasslands may be either overstated or understated. The same problem extends to carbon accounting. ANPP underpins estimates of carbon inputs, biomass turnover, and ecosystem carbon balance, so bias in ANPP propagates directly into assessments of grassland carbon sinks and sources. It also extends to remote sensing. Ground-based ANPP measurements are commonly treated as truth for calibrating or validating spatial products, yet those data may already embed OAC or OGC bias, allowing methodological error to be transferred into satellite-derived estimates and predictive models^38–42^.

The challenge is especially acute in grazed landscapes where recovery is already constrained. Restoration theory suggests that once chronic overgrazing, altered disturbance regimes, or land conversion erode belowground structure and recovery potential, undercompensation becomes more likely and enclosure-based estimates become more prone to overestimating ANPP. This risk is unlikely to be captured fully by existing datasets, particularly because our analyses include only domestic livestock and not the potentially substantial effects of large wild herbivores or invertebrate herbivory^43,44^. More generally, grassy ecosystems are regulated by interacting feedbacks among herbivory, fire, climate, and vegetation history^3,20^, so no single universal correction is likely to apply across grassland types and management systems. Improving ANPP estimation in grazed grasslands will therefore require not only better models, but field measurements that explicitly account for both biomass removal and the conditions governing post-grazing recovery.

## Methods

### Grazing experiment at Haibei research station

Since 2019, we have conducted a grazing experiment at the Haibei Alpine Grassland Ecosystem Research Station (Haibei research station), located in the north-eastern Tibetan Plateau (Fig. 1b). This region is influenced by a continental monsoon climate, featuring long, cold winters and short, cool summers. Over the past three decades, the average annual temperature has been −1.1 °C, with an average annual precipitation of 487.8 mm, approximately 84% of which occurs during the growing season from May to September^45,46^. These grasslands, historically subjected to moderate grazing pressure by livestock primarily during the growing season, have shown no signs of degradation.

#### Experiment design

We selected two flat, homogeneous areas as grazing sites (Site A and Site B), each covering an area of 150 m × 250 m and subjected to identical experimental treatments. Within each site, we implemented a randomized block design consisting of three blocks, with each block assigned four levels of grazing intensity. The grazing intensities were established according to the carrying capacity standards for natural grasslands outlined in Calculation of Rangeland Carrying Capacity Standard (NY/T635-2015). The experimental treatments include ungrazing, light grazing (0.5 yaks per hectare), moderate grazing (1.0 yak per hectare), and heavy grazing (2.0 yaks per hectare). Each experimental plot measures 40 m × 50 m, and barbed wire fences were installed between the plots. Grazing treatments were conducted using a rotational approach, with each block grazing for two days each fortnight.

#### ANPP monitoring

For each ungrazing plot, ANPP was monitored in mid-August, when plant growth typically reaches its peak, by weighing the dry matter of aboveground living biomass in five 0.5 m × 0.5 m quadrats. For each grazing plot, ANPP was calculated as the sum of the peak standing biomass and the biomass consumed by yaks prior to the peak standing biomass measurement. Biomass consumption at each grazing event was quantified by measuring the biomass differences inside and outside of the cages, and the total animal consumption was calculated by summing the consumptions of all grazing event (Supplementary Fig. 1).

For ungrazing plots,

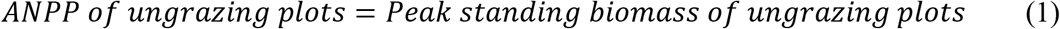

For grazing plots,

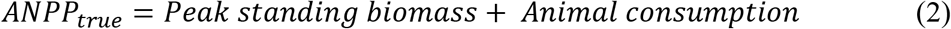

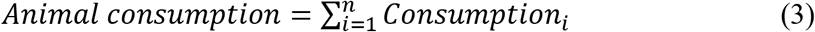

where ANPP_true_ is the true ANPP of grazing grassland (hereafter). Consumption is the animal consumption at each grazing event (Supplementary Figs. 2 and 3), i is the counts of grazing event.

#### Calculation of ANPP estimation errors

The ‘peak standing biomass’ approach in field surveys typically estimates ANPP_true_ by measuring aboveground living biomass during the peak growing season, either in grazing where animal consumption is not accounted for (Overlooked animal consumption, OAC method) or in ungrazing areas where plant growth compensation is overlooked (Overlooked growth compensation, OGC method).

Based on Equations (1-3), we further calculated the estimation errors of both methods using experimental observation data.

For OAC, the error was calculated as:

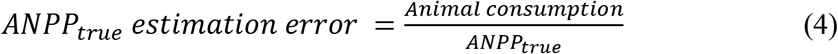

For OGC, the error was defined as:

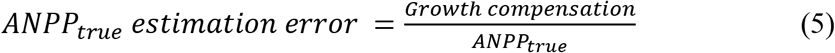

where growth compensation was quantified as:

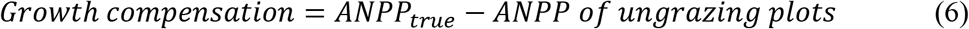

#### Statistical analyses

We conducted linear mixed models to analyze the effects of grazing intensity treatment, experimental sites (Site A and Site B), and observation year (2021–2023) on ANPP_true_ and its components (peak standing biomass, animal consumption and growth compensation). In the models, grazing treatment, site, year, and their interactions were treated as fixed effects, while block was included as a random effect to account for spatial variability among experimental blocks. The model was fitted using the *mixedlm* function in Python (via the *statsmodels* package), with parameter estimation conducted using maximum likelihood.

### Meta-analysis of Chinese grassland grazing experiments

#### Data collection

We collected peer-reviewed literature on the effects of grazing on ANPP using the Web of Science and China National Knowledge Infrastructure search engines (Supplementary Fig. 4). We identified relevant studies using the following keywords: “TS = ((Graz* OR herbivory OR defoliation) AND (plant OR vegetation & forage) AND (biomass OR productivity OR production OR ANPP OR NPP) AND (grassland OR steppe OR meadow OR pasture OR rangeland))”. The search results were filtered to include articles published between 1990 and 2024. Peer-reviewed studies were filtered based on the following criteria: (1) the intensity of grazing was defined in quantitative or qualitative terms; (2) the number of livestock foraging days was recorded; (3) climate conditions, soil properties, and species composition were similar in both grazing and ungrazing plots prior to grazing treatments; and (4) the mean, standard deviation/error, and sample size of ANPP or its components (standing biomass, ANPP, animal consumption, or/and growth compensation) can be directly extracted from the main text, tables, or figures. Details of the preferred items from systematic reviews are represented in Supplementary Fig. 4. Overall, 152 peer-reviewed articles met these criteria (Supplementary Table 8, nos. 1–152). Combined with our experimental data from the Haibei Research Station, this yielded a meta-analysis dataset for the China region comprised 936 individual observations, of which 217 were from ungrazed plots and 719 from grazed plots.

We also collected the information on latitude, longitude, grassland type, livestock species, grazing intensity (classified as ungrazing, light, moderate and heavy grazing, corresponding to grazing rates of 0, 0-30%, 30-60% and >60%, respectively^32,47^, foraging days, the size of livestock population, and grazing regime (continuous grazing or rotational grazing) for each selected paper.

#### Meta-analysis

For each grazed plot observation, we calculated the ANPP estimation error using the OAC method (Equation 4). For studies without reported animal consumption (Supplementary Table 8, nos. 1–120), consumption was estimated as:

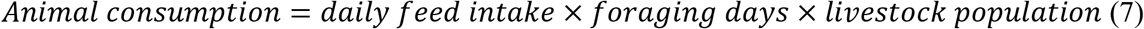

where daily feed intake was defined as 1.8 kg/day for a standardized sheep unit, following the Calculation of Rangeland Carrying Capacity Standard (NY/T635-2015)^48^.

For paired datasets consisting of both grazing exclusion and grazing intensity treatments, ANPP estimation errors were calculated using the OGC method (Equation 5). When studies reported multiple grazing intensity, each intensity was paired with its corresponding ungrazed treatment, yielding two or more ANPP estimation errors per study.

These case-specific ANPP estimation errors were synthesized using hierarchical mixed-effects meta-analysis implemented with the “rma.mv” function in the R package “metafor”. This analysis included the variable “Study ID” as a random factor to account for studies that contained multiple cases (i.e., more than one ANPP estimation value) and employed a weighting factor that assigned greater importance to cases with higher precision estimates. The weight factor was calculate using the sample size (N) (Equation 8):

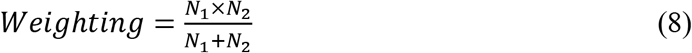

For the OAC method, N_1_ and N_2_ were assigned the same sample size corresponding to the grazing treatment. For the OGC method, N_1_ and N_2_ represent the sample sizes for the ungrazing and grazing treatments, respectively.

#### Analysis of drivers

We used the Q statistics to assessed the heterogeneity of the ANPP estimation error among studies^49^. Total heterogeneity (Q_T_) was partitioned into heterogeneity explained by moderator variables (Q_M_) and residual heterogeneity within groups (Q_R_). The moderators examined included grazing intensity, grazing regime, and climate zones characterized by temperature and moisture. Grazing intensity (light, moderate, heavy) and regime (continuous, rotational) were derived from the experimental settings of each study. Temperature zones were categorized as alpine, temperate, or tropical following the World Wide Fund for Nature’s ecoregion classification (https://ecoregions.appspot.com)^50^. Moisture zones were classified using the Standardized Precipitation-Evapotranspiration Index (SPEI), where SPEI≤-0.5 denotes arid, -0.5<SPEI<0.5 indicates semi-arid, and SPEI≥0.5 indicates humid conditions^51^. To quantify the explanatory power of each moderator, we calculated R²mod, defined as the proportion of total heterogeneity explained by the moderator.

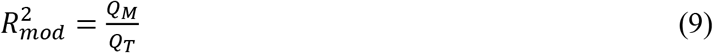

### Global assessment of ANPP estimation errors

We evaluated the estimation error of ANPP_true_ induced by grazing across the whole global grasslands and five main grassland ecoregions —North American Prairie, Eurasian Steppe, Alpine grassland, South American pampas, Savanna, which are classified by climate type and vegetation structure according to global meta-data distribution^50^.

#### Estimation error from OAC

ANPP estimation error under the OAC method was estimated from multi-year averaged global spatial raster datasets of peak standing biomass and animal consumption (Supplementary Fig. 5).

Peak standing biomass data are obtained from a publicly available dataset provided by Oak Ridge National Laboratory Distributed Active Archive Centre^52^. Animal consumption was estimated based on livestock numbers, daily feed intake and foraging days (Equation 7). Livestock numbers were obtained from the Gridded Livestock of the World (GLW) database produced by the Food and Agriculture Organization (FAO) (https://data.apps.fao.org). We used an averaged livestock density map derived from three datasets: GLW3-2010, GLW4-2015, and GLW4-2020. Livestock numbers were conversed to sheep unit (SU)^37^, and only populations within areas classified as natural grasslands were included^53^. Foraging days were defined as the period from the start of growing season to the peak of the growing season, and it is derived from the global land surface phenology (GLSP) dataset (https://www.earthdata.nasa.gov). Details of all the data sources are displayed in Supplementary Table 6.

#### Estimation error from OGC

Due to the lack of the spatial data of compensation effects, point data of this effects were derived for five grassland ecoregions based on meta-analysis. For consistency and comparability, only observations corresponding to moderate grazing intensity were extracted from each study, based on the assumption that most grasslands globally experience moderate grazing pressure. Thus, ANPP estimation error under the OGC approach was derived from the moderate grazing treatment based on a global meta-analysis. Building on the criteria established for the meta-analysis of Chinese grasslands, additional standards were applied for the global assessment to ensure methodological consistency and data quality. Because rotational grazing studies remain scarce at the global scale, the present analysis was confined to continuous grazing systems. To improve the accuracy of ANPP estimation, only studies that directly reported either true ANPP or animal consumption were included. The final meta-analysis dataset comprised 58 studies and 212 observations (Supplementary Table 8, nos. 121–178; Supplementary Fig. 4). Among these, 32 studies from China (Supplementary Table 8, nos. 121–152) providing either true ANPP or animal consumption data were used for global prediction.

Growth compensation factor (F) was introduced to represent the compensation effects (Supplementary Table 7). It represents the difference in ANPP between moderately grazing and ungrazing treatments, was calculated as:

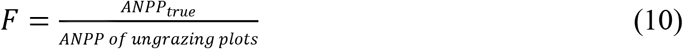

where F < 1 indicates under-compensation, whereas F > 1 denotes over-compensation. According to equations 5, 6 and 10, ANPP_true_ estimation error can be calculated as:

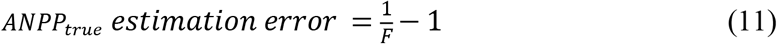

As the OGC results were available as point data, while the OAC results as raster data, it was necessary to ensure spatial correspondence between the two estimation approaches. To achieve this, the OAC values were extracted for the same locations as the OGC points by averaging the surrounding 10 × 10 raster pixels (1 km) for each point.

## Data availability

All data are available in the Supplementary Information.

## Acknowledgements

We are grateful for the constructive comments provided by Zhenong Jin, Yanhong Tang, and David J Eldridge on early versions of this manuscript. This work was supported by the National Natural Science Foundation of China (Grants No. 32192461, 32130065, 32422058), the Major Project of Science and Technology Development in Gansu Province (23ZDNA009), and the Postdoctoral Fellowship Program of CPSF (GZC20230998).

## Author contributions

Conceptualization: J.-S.H.; Methodology: Y.S., H.W., H.Z., J.-S.H.; Investigation: H.Z., X.W., Y.S., J.-S.H.; Visualization: Y.S., X.W.; Funding acquisition: J.-S.H.; Project administration: J.-S.H.; Supervision: J.-S.H.; Writing – original draft: Y.S., H.W., J.-S.H.; Writing – review and editing: All co-authors.

## Supplementary Information

**Supplementary Figure 1.**
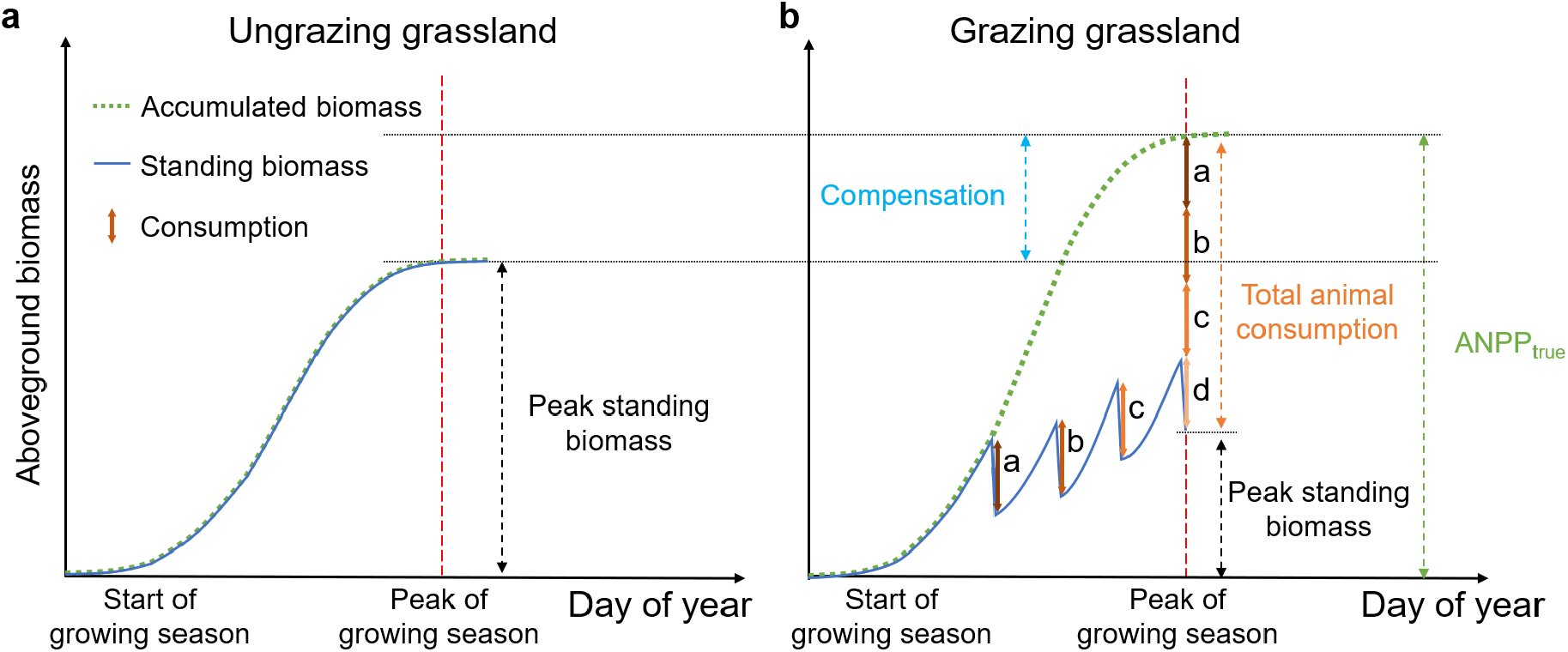
Schematic illustration of aboveground biomass dynamics. **a**, Ungrazing grassland. **b**, Grazing grassland. Letters a, b, c and d represent the biomass consumed by livestock during each grazing event. ANPP_true_ is the true value of ANPP for grazing grassland.

**Supplementary Figure 2.**
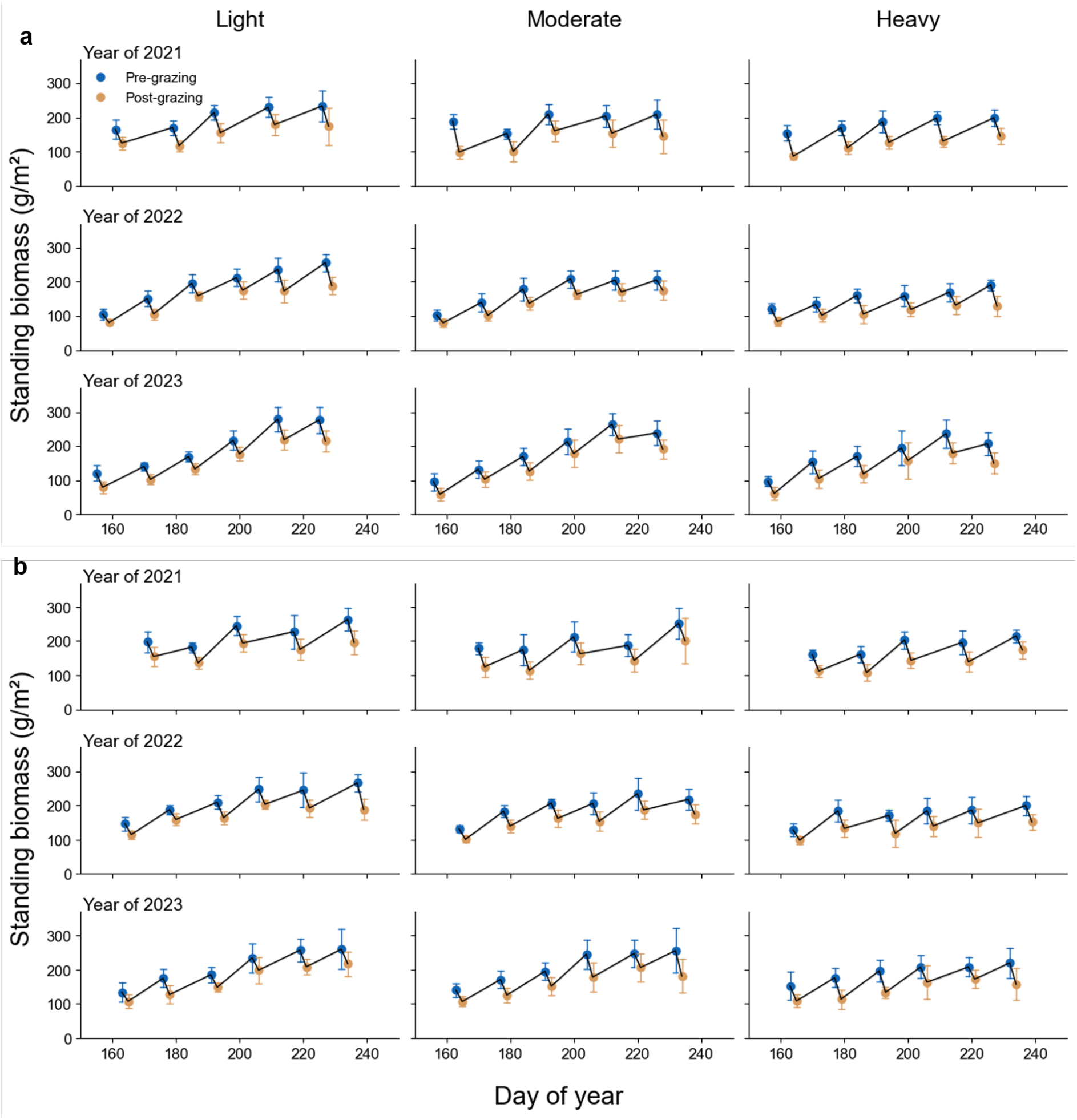
Standing biomass of pre- and post-grazing from 2021 to 2023. **a**, Standing biomass from experiment site a. **b**, Standing biomass from experiment site b.

**Supplementary Figure 3.**
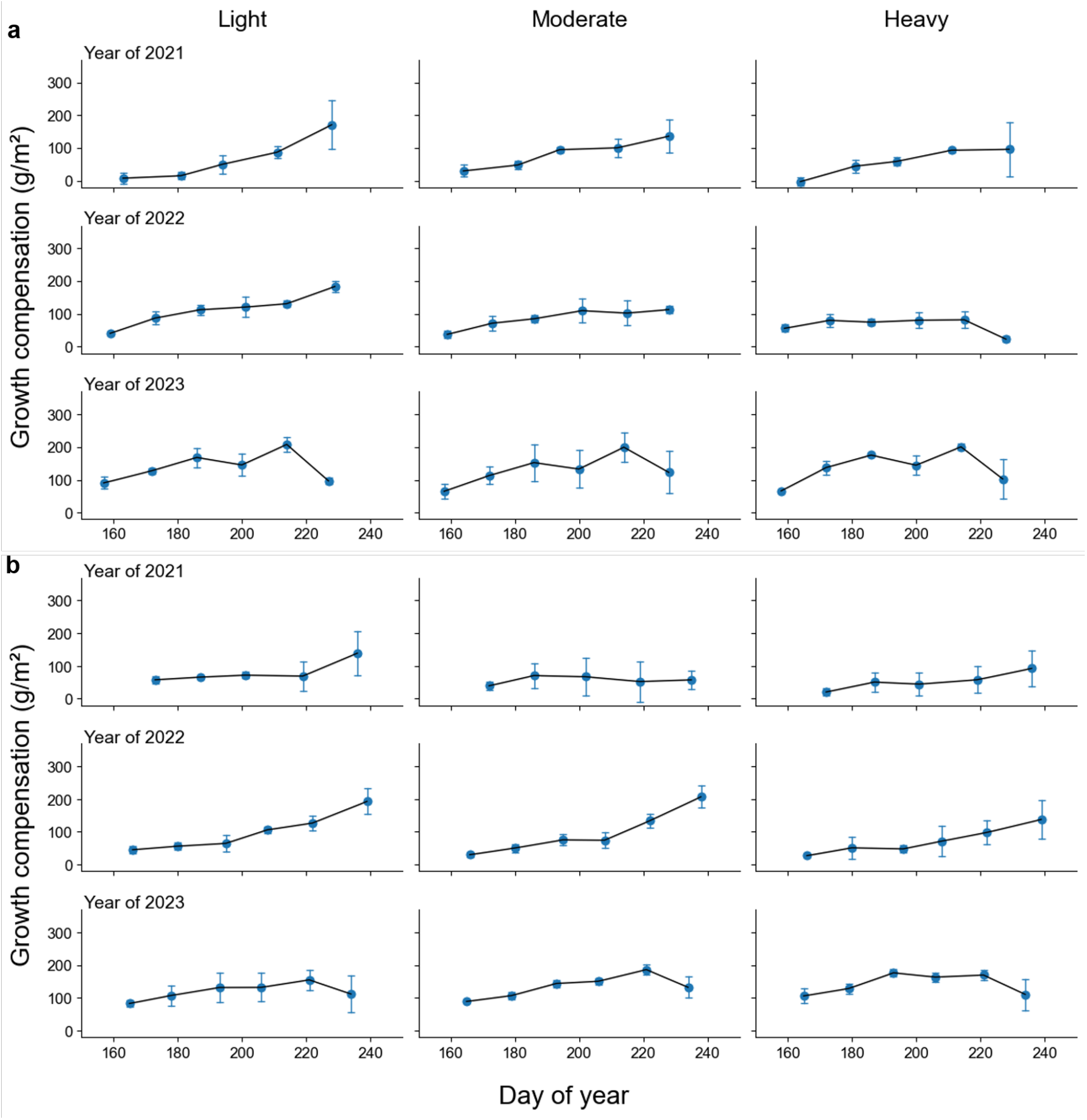
Growth compensation at each grazing event from 2021 to 2023. **a**, Growth compensation from experiment site a. **b**, Growth compensation from experiment site b.

**Supplementary Figure 4.**
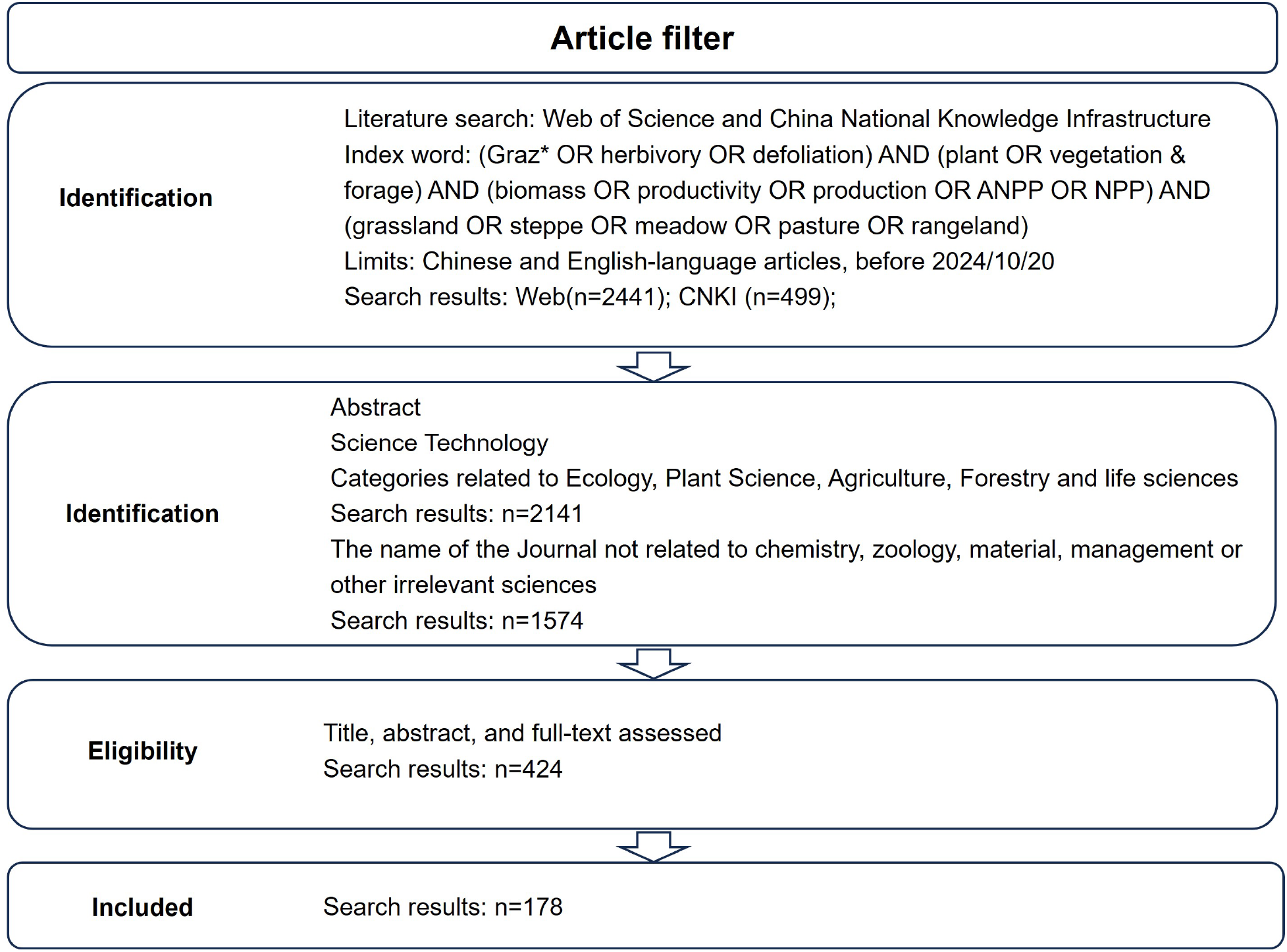
Processes of filtering literature for grazing effects on aboveground net primary productivity following the guidelines of the preferred items for systematic reviews and meta-analyses of Chinese grassland (PRISMA).

**Supplementary Figure 5.**
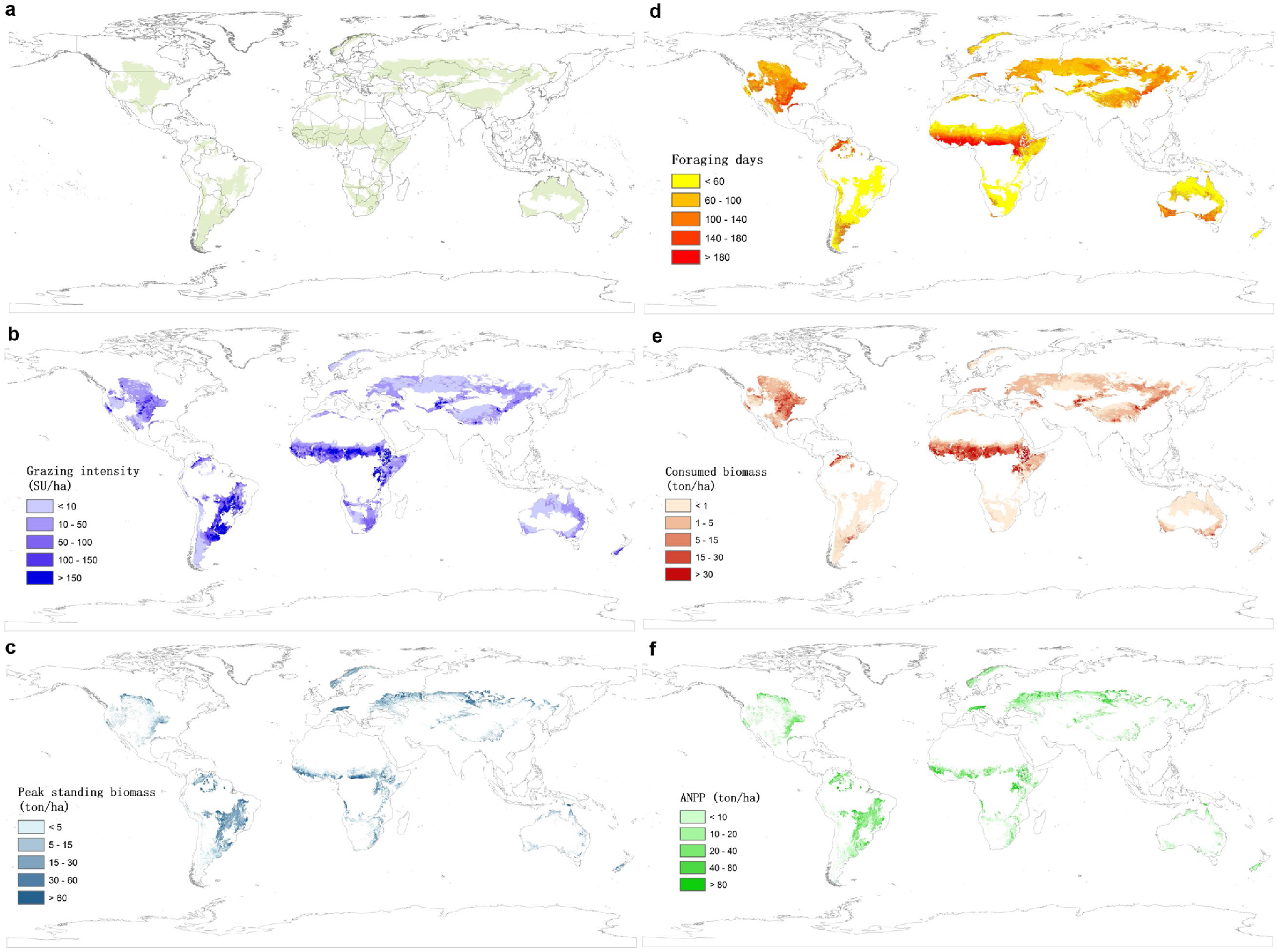
Global patterns of metrics related to grazing and aboveground net primary production (ANPP) in natural grasslands. **a**, Grassland areas. **b**, Grazing intensity. **c**, Peak standing biomass. **d**, Foraging days. **e**, Consumed biomass. **f**, ANPP.

**Supplementary Table 1.** Summary of ANPP and its components across treatments in the alpine grassland experiment.

| **Treatment** | **2021** | **2022** | **2023** | **2021-2023** |
| --- | --- | --- | --- | --- |
| Peak standing biomass |  |  |  |  |
| Ungrazing | 348.79a | 312.38a | 328.45a | 329.87a |
| Light | 233.27b | 245.80b | 278.00a | 252.36b |
| Moderate | 213.45bc | 229.30bc | 252.98ab | 231.91bc |
| Heavy | 164.05d | 171.75d | 165.66c | 167.15d |
| Animal consumption |  |  |  |  |
| Ungrazing | - | - | - | - |
| Light | 270.61a | 255.00a | 135.72b | 220.44a |
| Moderate | 232.16a | 243.26a | 205.58b | 227.00a |
| Heavy | 279.02a | 221.04a | 272.00a | 257.35a |
| Growth compensation |  |  |  |  |
| Ungrazing | - | - | - | - |
| Light | 155.09a | 188.41a | 82.55b | 142.02a |
| Moderate | 96.81b | 160.16a | 127.38a | 128.12ab |
| Heavy | 94.28b | 80.40b | 106.48a | 93.72b |
| ANPP |  |  |  |  |
| Ungrazing | 348.79b | 312.38c | 328.45b | 329.87b |
| Light | 503.88a | 500.80a | 413.72a | 472.80a |
| Moderate | 445.60a | 472.54a | 458.56a | 458.90a |
| Heavy | 443.07a | 392.79b | 437.65a | 424.50a |

#Unit: g/m<sup>2</sup>. Lowercase letters denote significant differences ( $p < 0.05$ ).

**Supplementary Table 2.** Statistical results of linear mixed-effects models examining the individual and interactive effects of treatment, site, and observation year in the grazing experiment.

|  | Site<br>(S) | Treatment<br>(T) | Year<br>(Y) | S * T | S * Y | T * Y | S * T *<br>Y |
| --- | --- | --- | --- | --- | --- | --- | --- |
| Peak standing biomass |  |  |  |  |  |  |  |
| *df* | 1 | 3 | 2 | 3 | 2 | 6 | 6 |
| *F* | 0.05 | 75.67 | 2.02 | 0.19 | 4.81 | 1.72 | 1.55 |
| *p* | 0.818 | < 0.001 | 0.144 | 0.900 | 0.012 | 0.137 | 0.182 |
| Animal consumption |  |  |  |  |  |  |  |
| *df* | 1 | 2 | 2 | 2 | 2 | 4 | 4 |
| *F* | 2.32 | 3.05 | 6.35 | 0.19 | 3.85 | 5.78 | 2.98 |
| *p* | 0.136 | 0.059 | 0.004 | 0.826 | 0.030 | 0.001 | 0.031 |
| Growth compensation |  |  |  |  |  |  |  |
| *df* | 1 | 2 | 2 | 2 | 2 | 4 | 4 |
| *F* | 1.64 | 3.48 | 2.13 | 0.42 | 4.36 | 2.79 | 1.05 |
| *p* | 0.208 | 0.041 | 0.133 | 0.660 | 0.020 | 0.041 | 0.393 |
| ANPP |  |  |  |  |  |  |  |
| *df* | 1 | 3 | 2 | 3 | 2 | 6 | 6 |
| *F* | 1.2 | 29.01 | 1.57 | 0.64 | 0.2 | 2.51 | 1.35 |
| *p* | 0.278 | < 0.001 | 0.218 | 0.596 | 0.820 | 0.034 | 0.253 |
| OAC (%) |  |  |  |  |  |  |  |
| *df* | 1 | 2 | 2 | 2 | 2 | 4 | 4 |
| *F* | 2.33 | 3.06 | 6.35 | 0.19 | 3.86 | 5.78 | 2.99 |
| *p* | 0.136 | 0.059 | 0.004 | 0.826 | 0.030 | 0.001 | 0.031 |
| OGC (%) |  |  |  |  |  |  |  |
| *df* | 1 | 2 | 2 | 2 | 2 | 4 | 4 |
| *F* | 1.65 | 3.49 | 2.13 | 0.42 | 4.36 | 2.79 | 1.06 |
| *p* | 0.208 | 0.041 | 0.133 | 0.660 | 0.020 | 0.041 | 0.393 |

#OAC, overlooked animal consumption; OGC, overlooked growth compensation.

**Supplementary Table 3.** Summary of the effects of grazing on ANPP estimation errors in the grazing experiment.

| **Treatment** | **2021** | **2022** | **2023** | **2021-2023** |
| --- | --- | --- | --- | --- |
| **OAC (%)** |  |  |  |  |
| Light | -53.02a | -50.82a | -32.01a | -45.28a |
| Moderate | -51.82ab | -51.68a | -44.44a | -49.31a |
| Heavy | -62.43b | -55.63a | -55.63b | -57.90b |
| **OGC (%)** |  |  |  |  |
| Light | -29.69a | -37.47b | -18.56a | -28.57b |
| Moderate | -20.83a | -33.27ab | -27.07a | -27.06ab |
| Heavy | -19.68a | -18.59a | -23.29a | -20.52a |

#OAC, overlooked animal consumption; OGC, overlooked growth compensation. Lowercase letters denote significant differences ( $p < 0.05$ ).

**Supplementary Table 4.** Summary of the effects of grazing intensities and regimes on ANPP estimation errors in the meta-analysis.

| **Factors** | **OAC (%)** | **OGC (%)** |
| --- | --- | --- |
| Overall | -38.32 | 11.99 |
| Grazing intensities |  |  |
| Light | -18.22a | 18.61a |
| Moderate | -40.51b | 19.72a |
| Heavy | -61.27c | 2.36b |
| Grazing regimes |  |  |
| Continuous grazing | -34.87a | 19.22a |
| Rotational grazing | -38.37a | -7.09b |
| Temperature zone |  |  |
| Alpine climate | -32.36a | 5.43a |
| Temperate climate | -47.54a | 15.52a |
| Moisture zone |  |  |
| Arid | -33.12a | 14.05a |
| Semi-arid | -49.45a | 8.61a |

#Positive values indicate overestimation, whereas negative values indicate underestimation.

**Supplementary Table 5.** Summary of the effects of grassland ecozones, temperature and moisture zones on ANPP estimation error in the global meta-analysis.

| **Factors** | **OAC (%)** | **OGC (%)** |
| --- | --- | --- |
| Overall | -34.25 | 14.46 |
| Grassland ecozones |  |  |
| North American grassland | -43.56c | 15.80a |
| Eurasian grassland | -32.69b | 14.62a |
| South American pampas | -33.97b | 21.41a |
| Savanna | -48.93d | 23.10a |
| Alpine grassland | -14.90a | 11.47a |
| Temperature zone |  |  |
| Alpine | -14.90a | 11.47a |
| Temperate | -36.23b | 15.11a |
| Tropical | -48.93bc | 23.10a |
| Moisture zone |  |  |
| Arid | -14.90a | 11.47a |
| Semi-arid | -38.80b | 15.12a |
| Humid | -41.58bc | 16.18a |

#OAC, overlooked animal consumption; OGC, overlooked growth compensation. Positive values indicate overestimation, whereas negative values indicate underestimation. Lowercase letters indicate significance ( $p < 0.05$ ).

**Supplementary Table 6.**
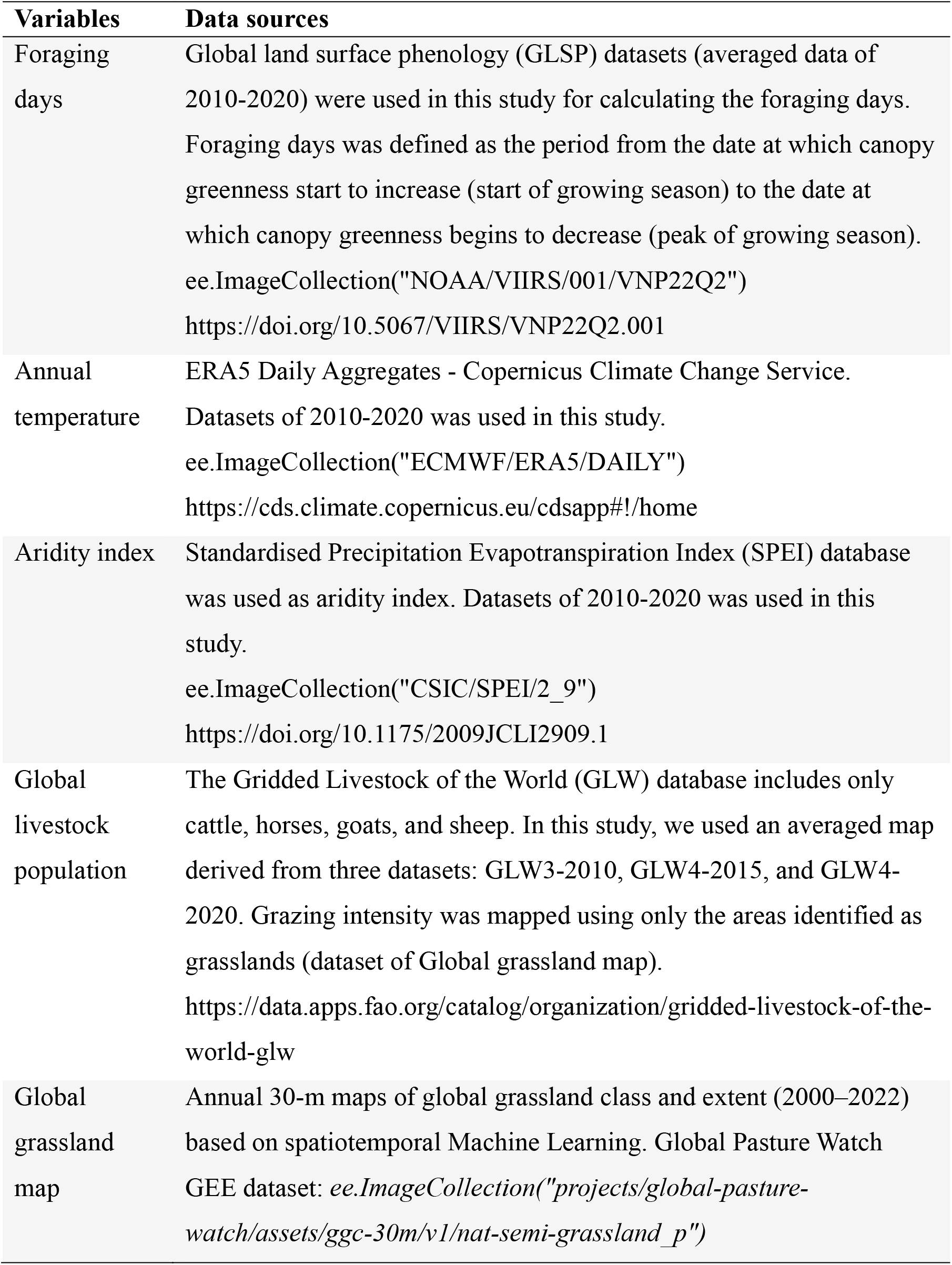
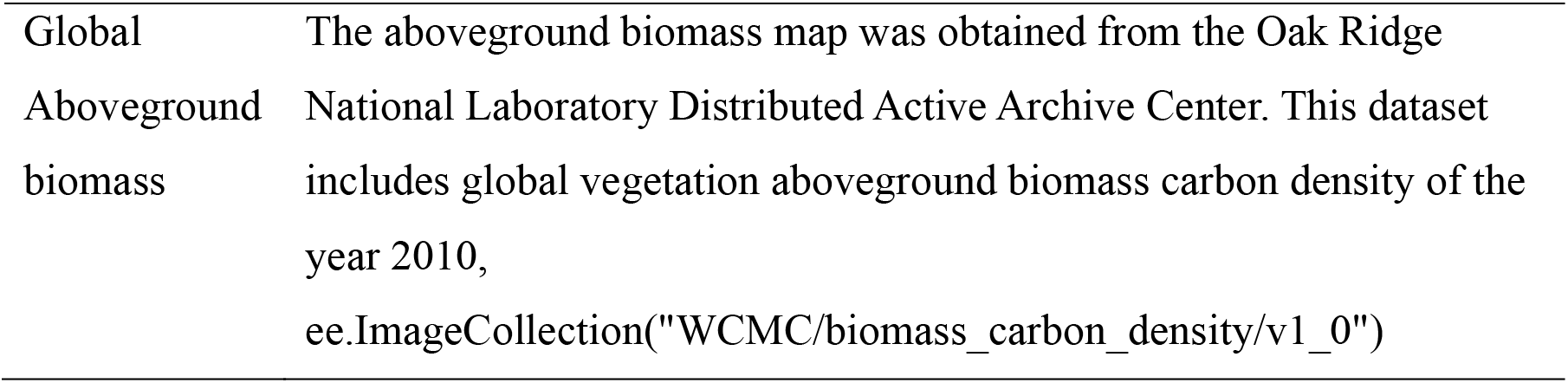
Data sources for the variables used.

| **Variables** | **Data sources** |
| --- | --- |
| Foraging days | Global land surface phenology (GLSP) datasets (averaged data of 2010-2020) were used in this study for calculating the foraging days. Foraging days was defined as the period from the date at which canopy greenness start to increase (start of growing season) to the date at which canopy greenness begins to decrease (peak of growing season).<br>ee.ImageCollection("NOAA/VIIRS/001/VNP22Q2")<br>https://doi.org/10.5067/VIIRS/VNP22Q2.001 |
| Annual temperature | ERA5 Daily Aggregates - Copernicus Climate Change Service. Datasets of 2010-2020 was used in this study.<br>ee.ImageCollection("ECMWF/ERA5/DAILY")<br>https://cds.climate.copernicus.eu/cdsapp#!/home |
| Aridity index | Standardised Precipitation Evapotranspiration Index (SPEI) database was used as aridity index. Datasets of 2010-2020 was used in this study.<br>ee.ImageCollection("CSIC/SPEI/2_9")<br>https://doi.org/10.1175/2009JCLI2909.1 |
| Global livestock population | The Gridded Livestock of the World (GLW) database includes only cattle, horses, goats, and sheep. In this study, we used an averaged map derived from three datasets: GLW3-2010, GLW4-2015, and GLW4-2020. Grazing intensity was mapped using only the areas identified as grasslands (dataset of Global grassland map).<br>https://data.apps.fao.org/catalog/organization/gridded-livestock-of-the-world-glw |
| Global grassland map | Annual 30-m maps of global grassland class and extent (2000–2022) based on spatiotemporal Machine Learning. Global Pasture Watch GEE dataset: *ee.ImageCollection("projects/global-pasture-watch/assets/ggc-30m/v1/nat-semi-grassland_p")* |
| Global<br>Aboveground<br>biomass | The aboveground biomass map was obtained from the Oak Ridge National Laboratory Distributed Active Archive Center. This dataset includes global vegetation aboveground biomass carbon density of the year 2010,<br>`ee.ImageCollection("WCMC/biomass_carbon_density/v1_0")` |

**Supplementary Table 7.** Information of growth compensation coefficient for five grassland ecozones.

| **Region** | **F** | **Observation No.** |
| --- | --- | --- |
| **Overall** | 0.936 | 203 |
| **North American Prairie** | 0.955 | 40 |
| **Eurasian Steppe** | 0.935 | 103 |
| **Alpine grassland** | 0.930 | 49 |
| **South American pampas** | 0.857 | 5 |
| **Savanna** | 0.914 | 6 |

#F is growth compensation coefficient. $F = \text{ANPP}_{\text{true}} / (\text{ANPP of ungazing plots})$ . $F < 1$ indicates under-compensation, and $F > 1$ indicates over-compensation.

**Supplementary Table 8.**
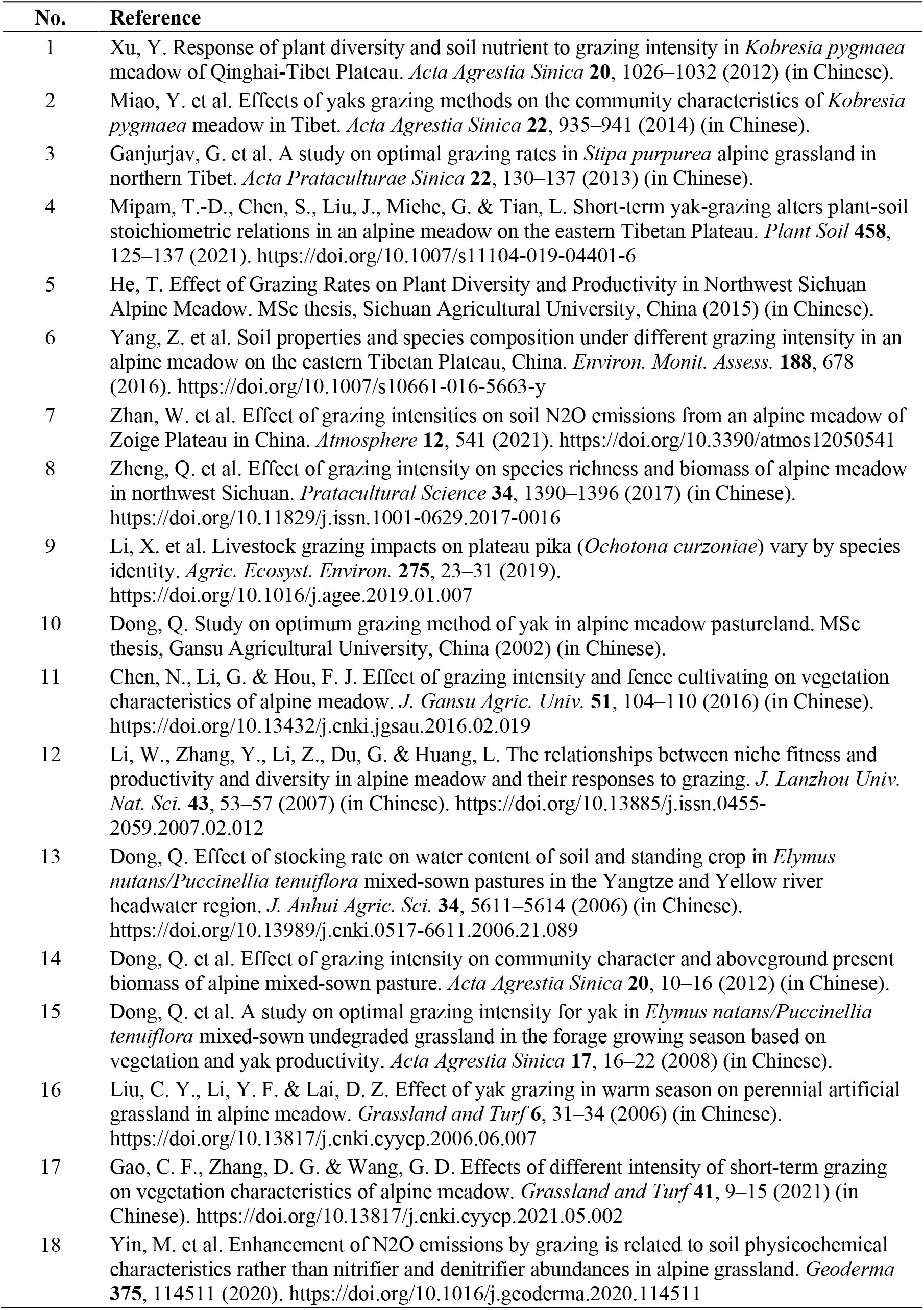
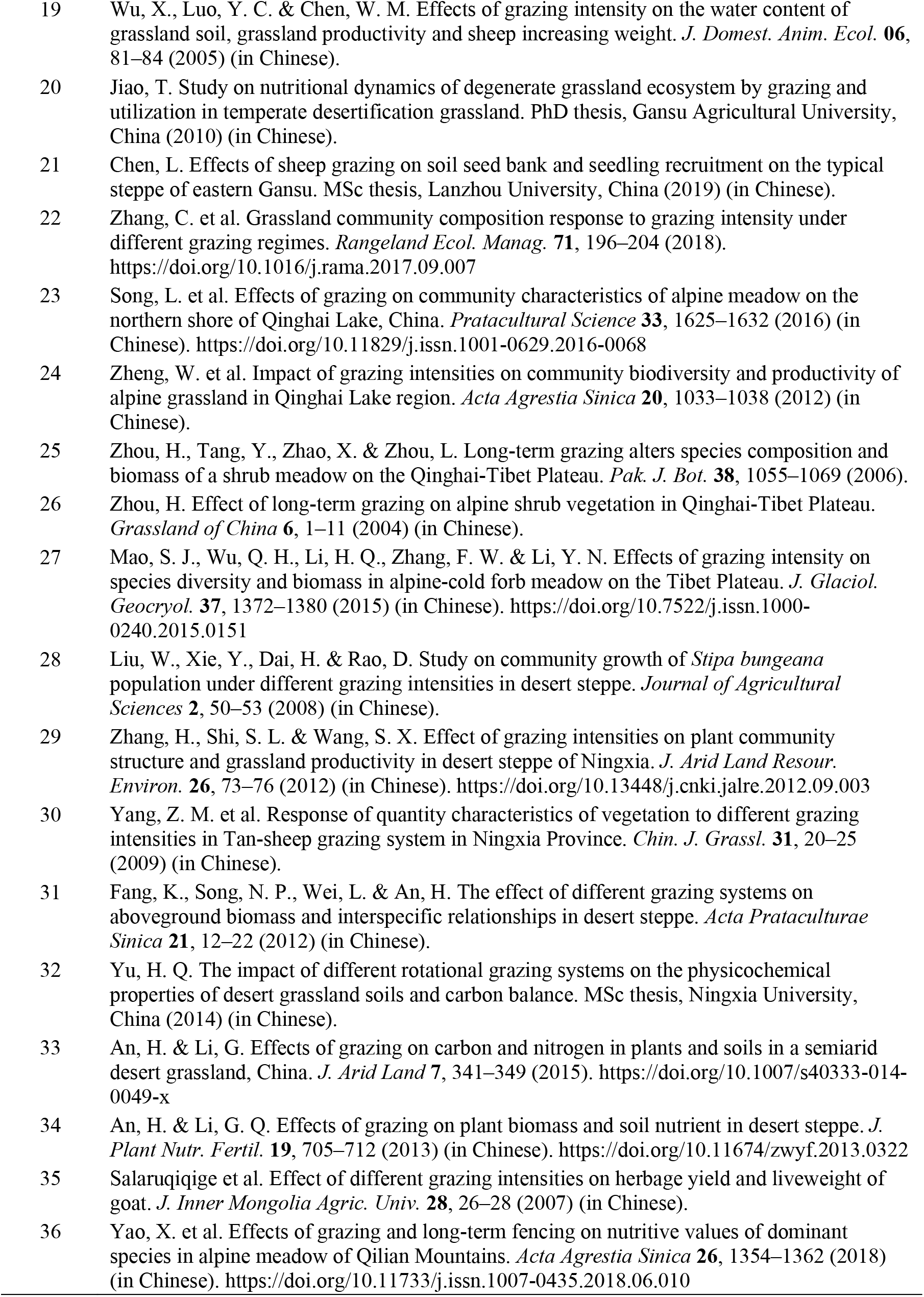
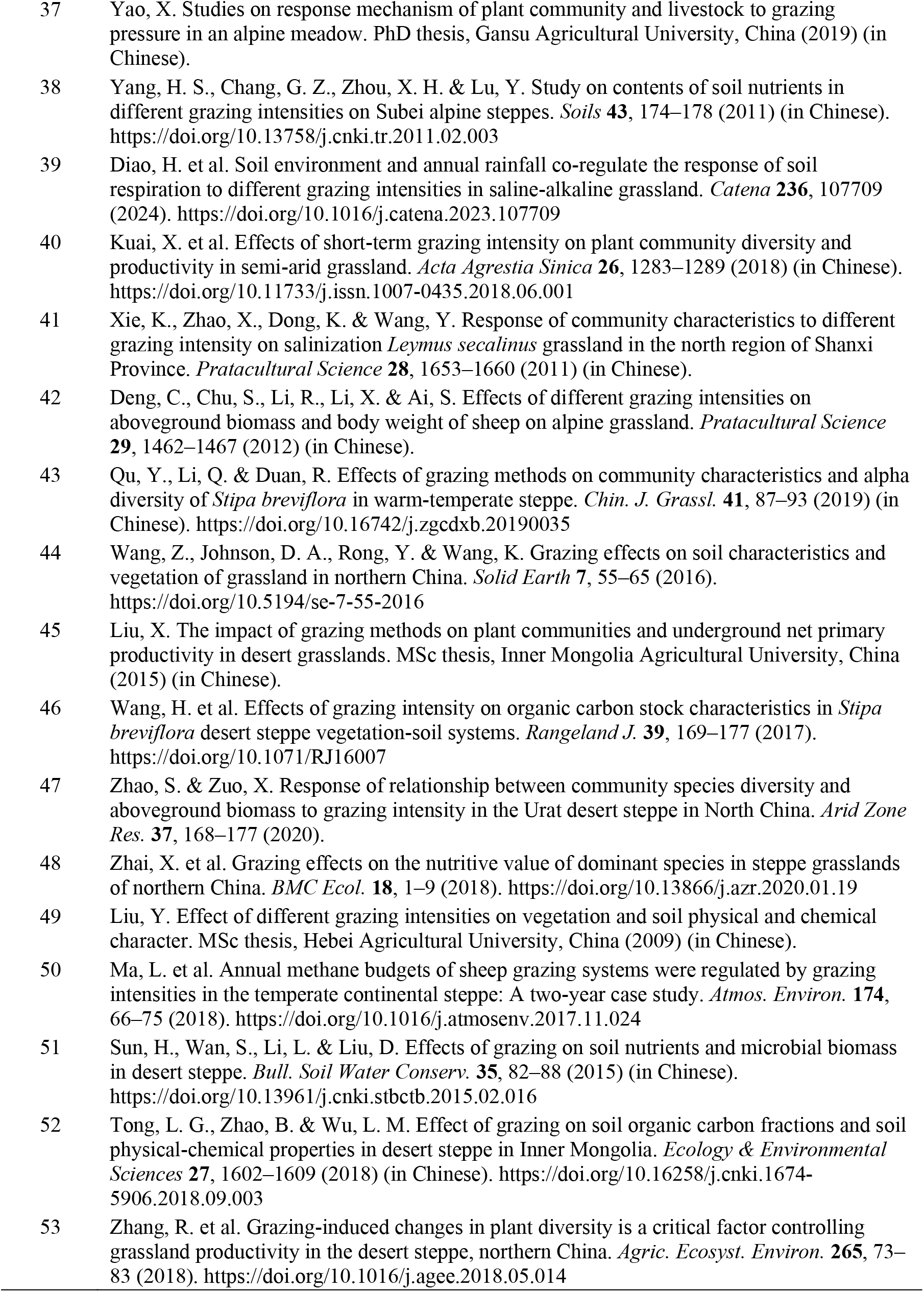
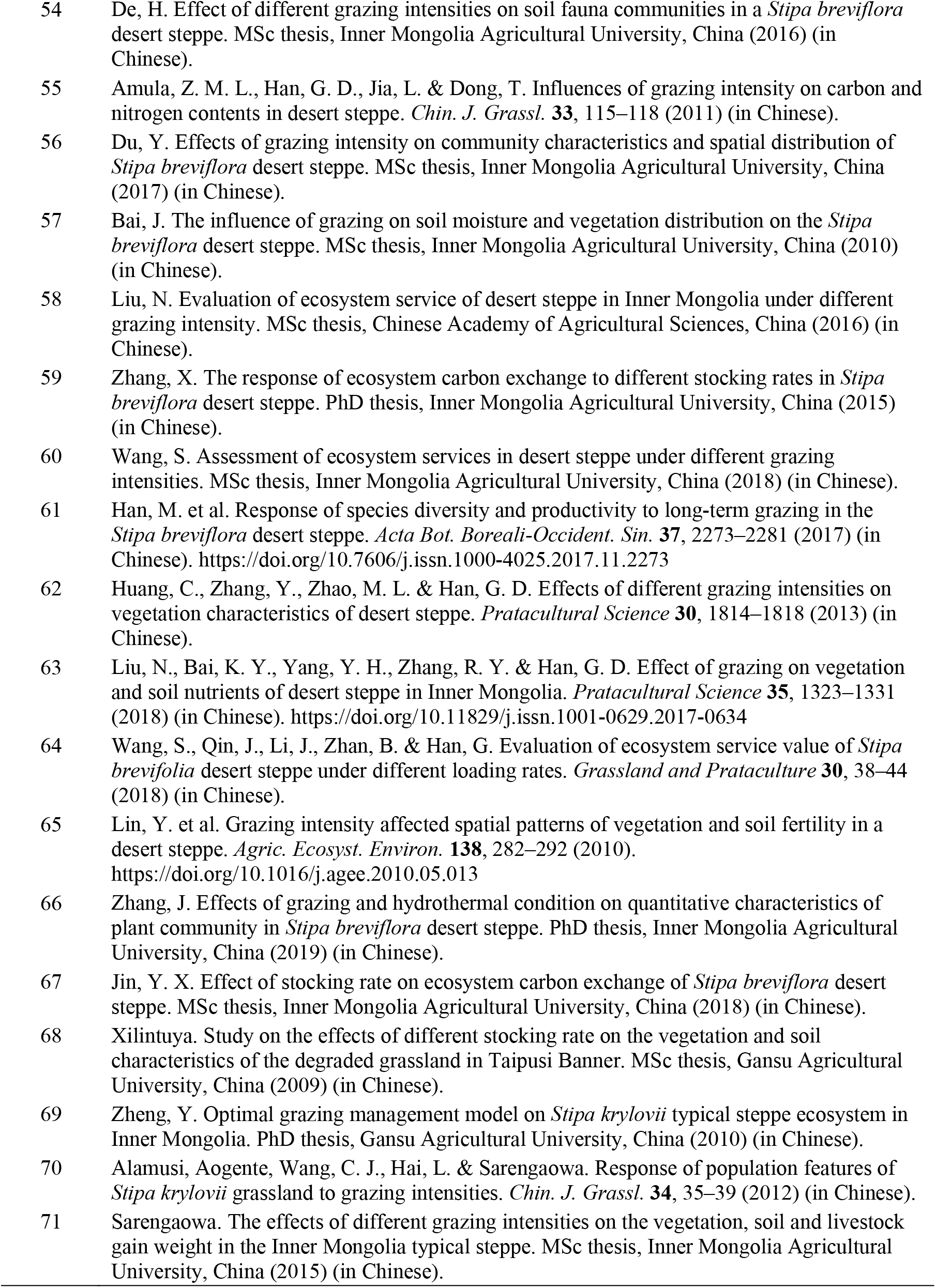
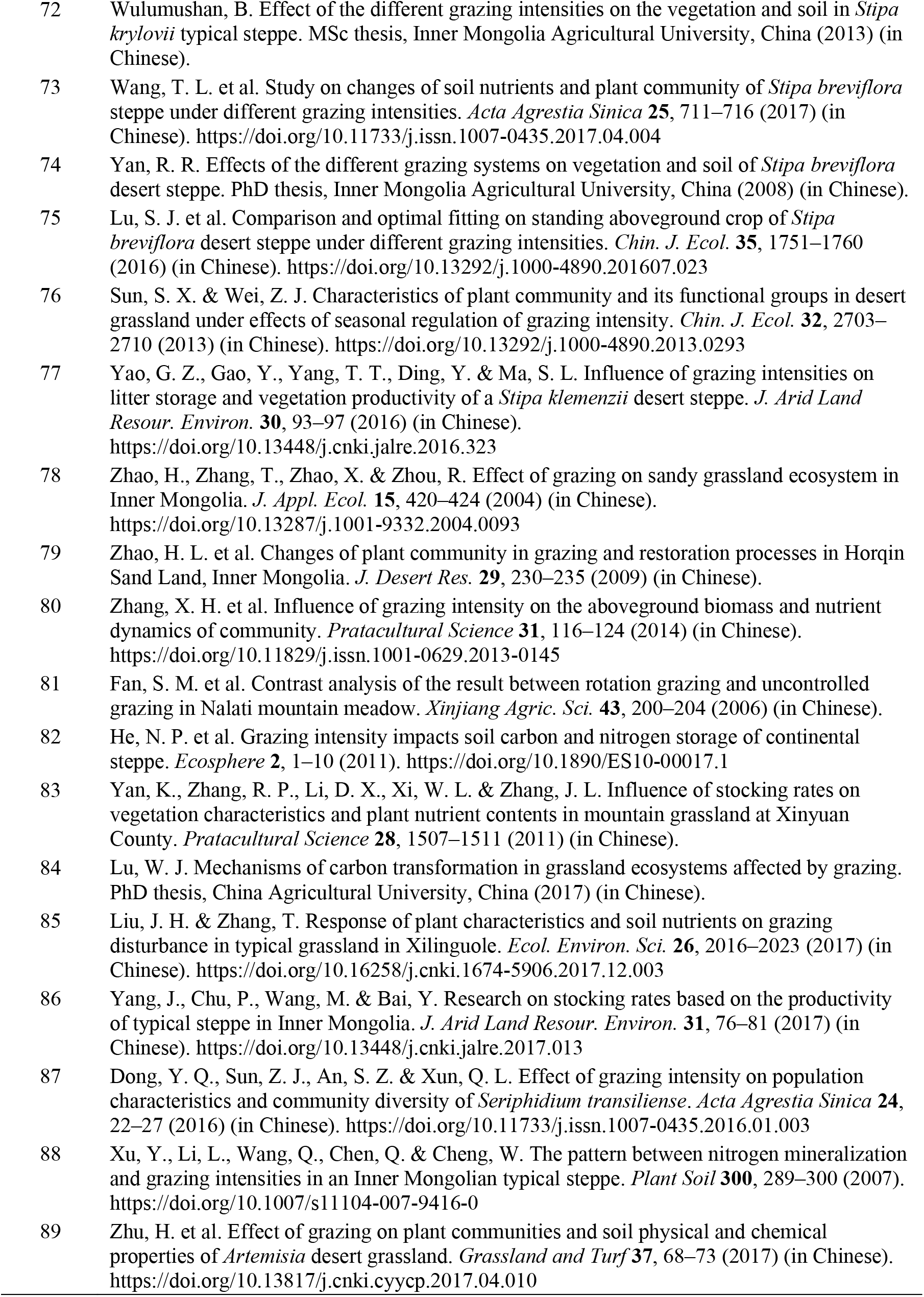
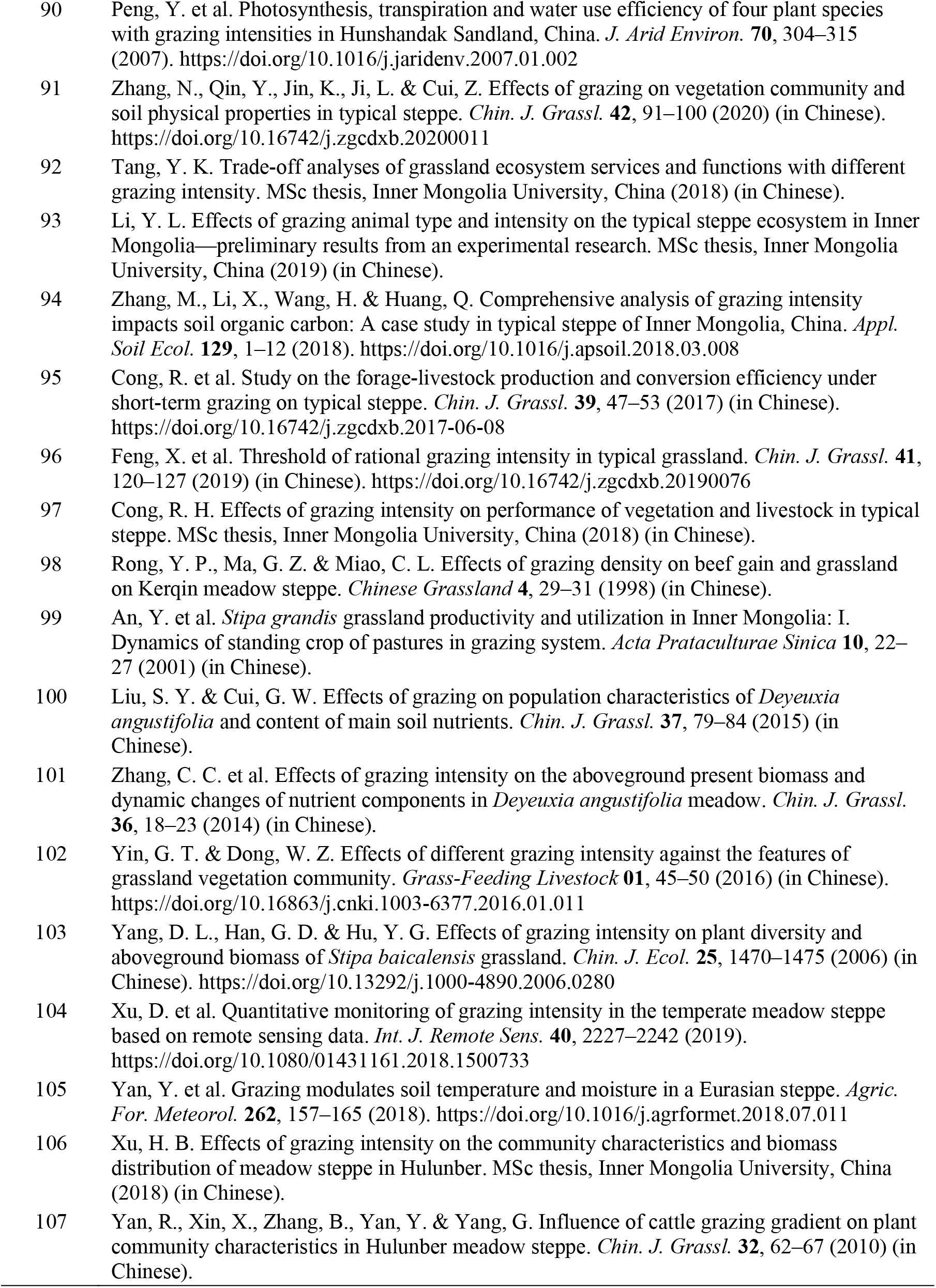
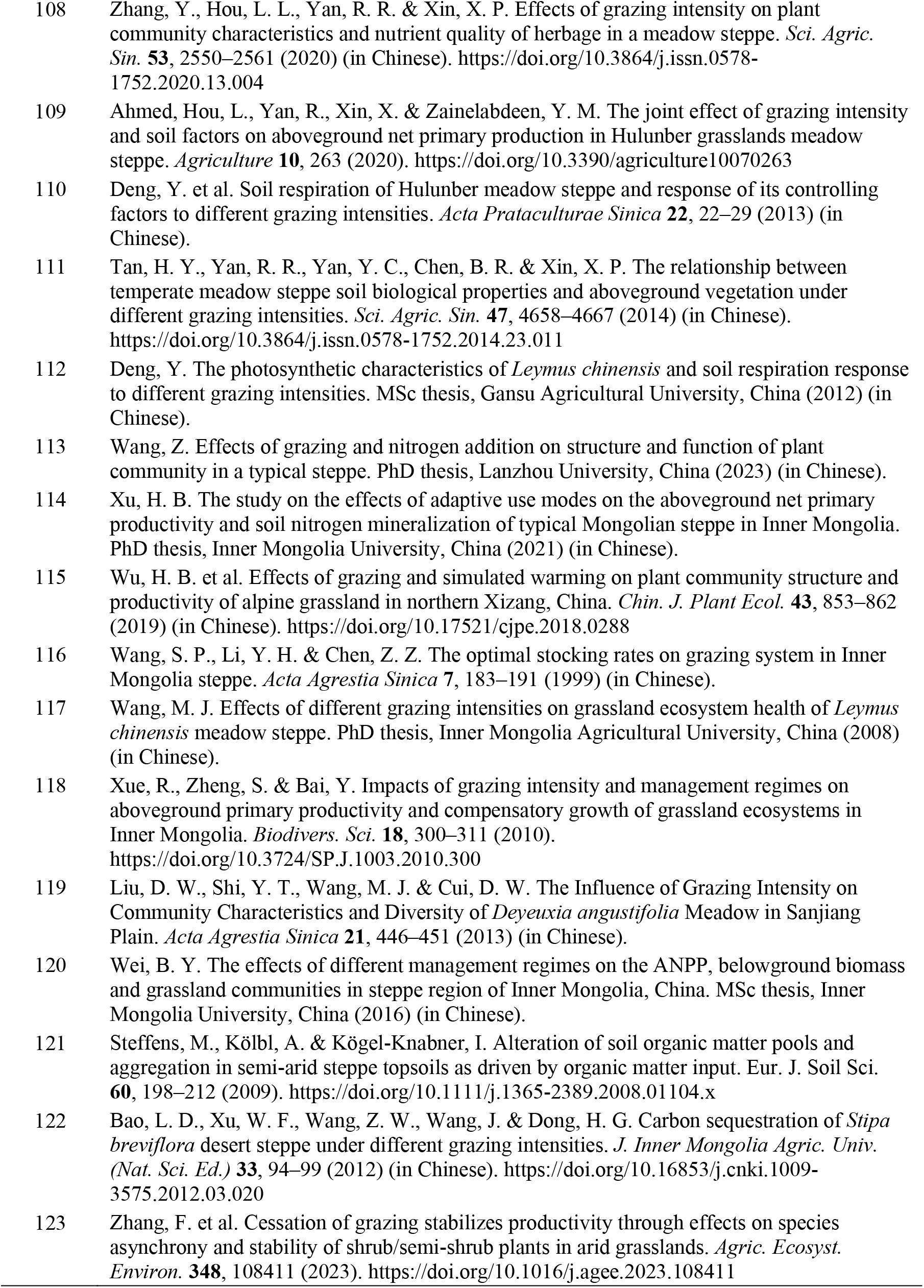
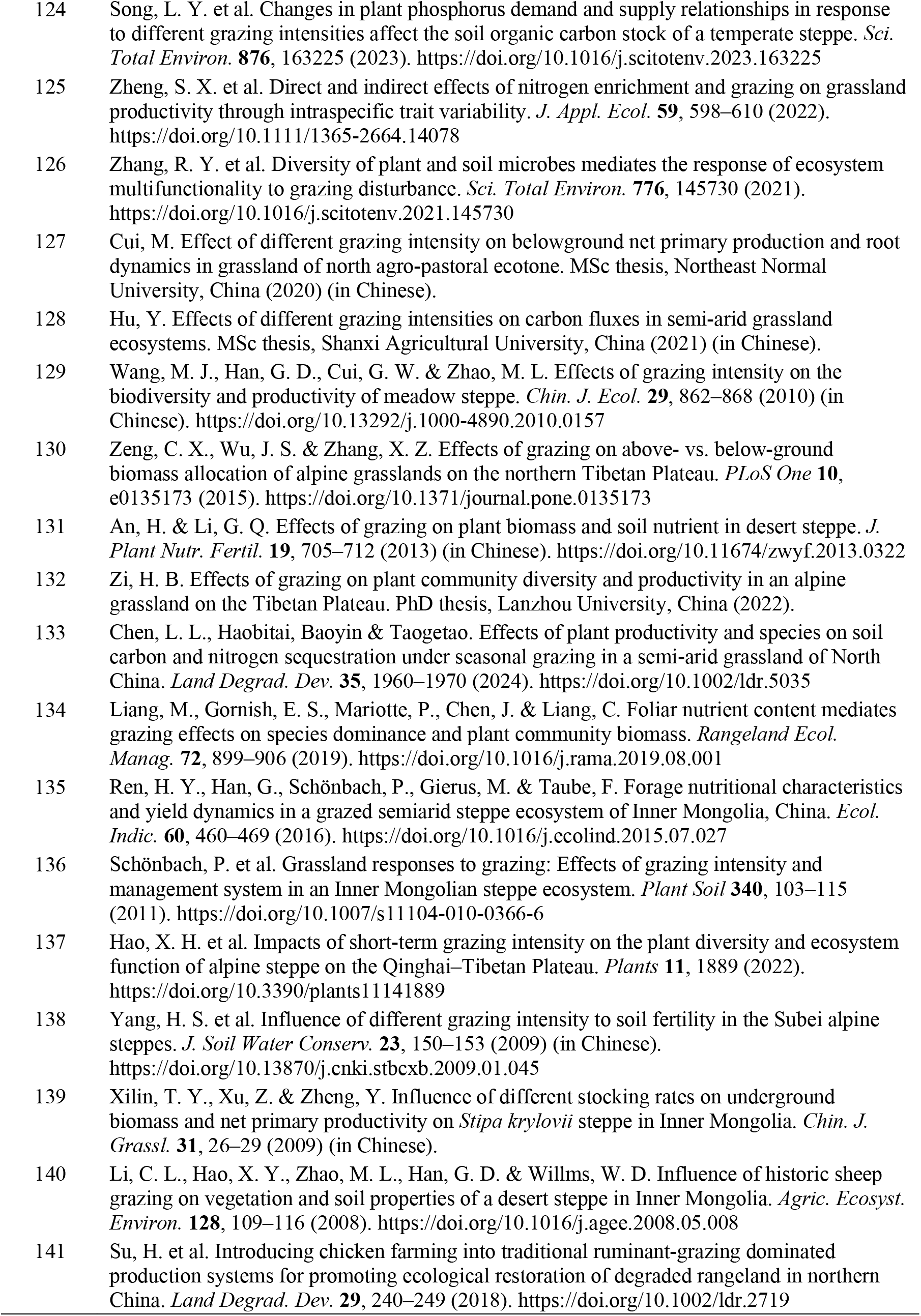
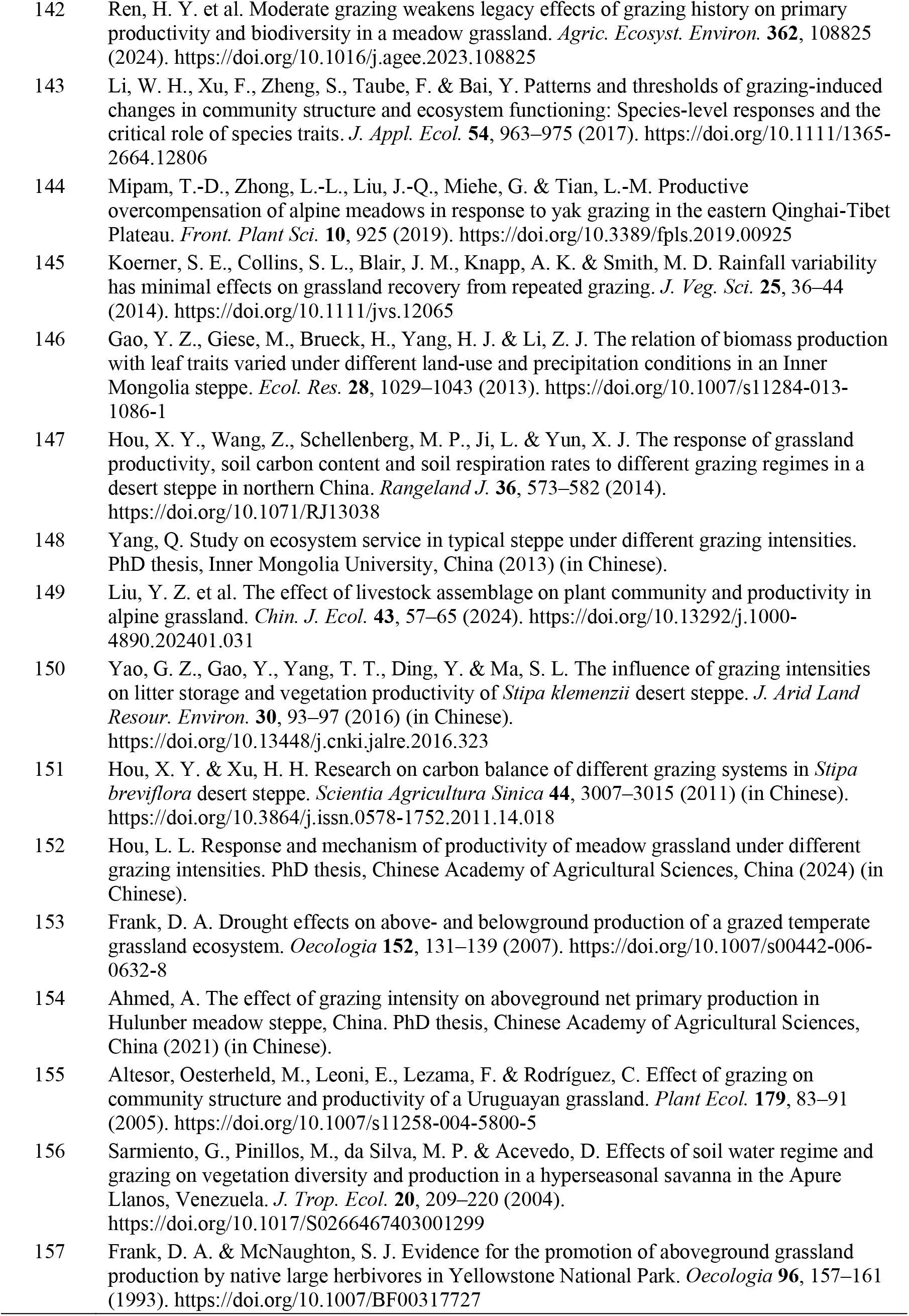
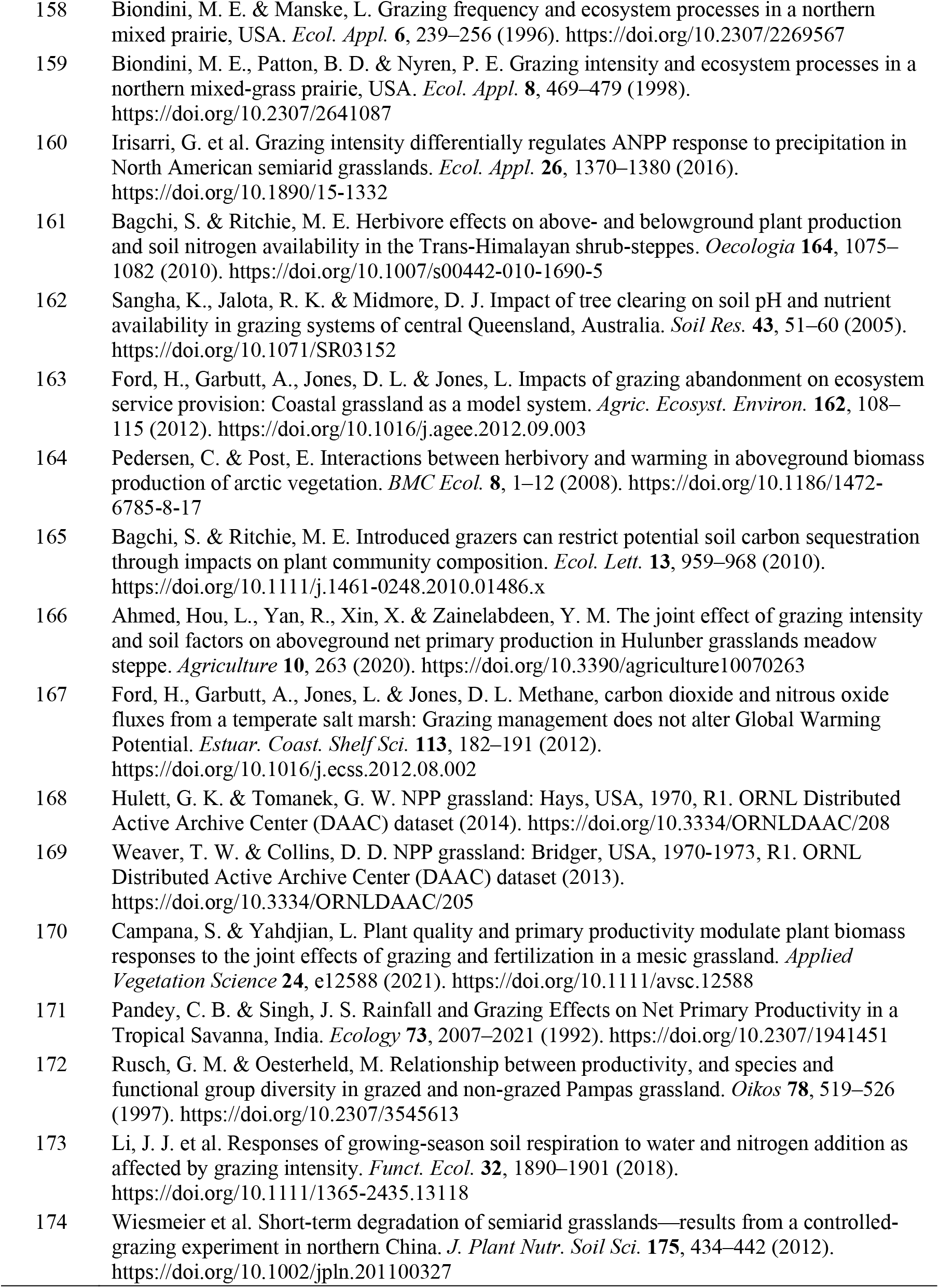
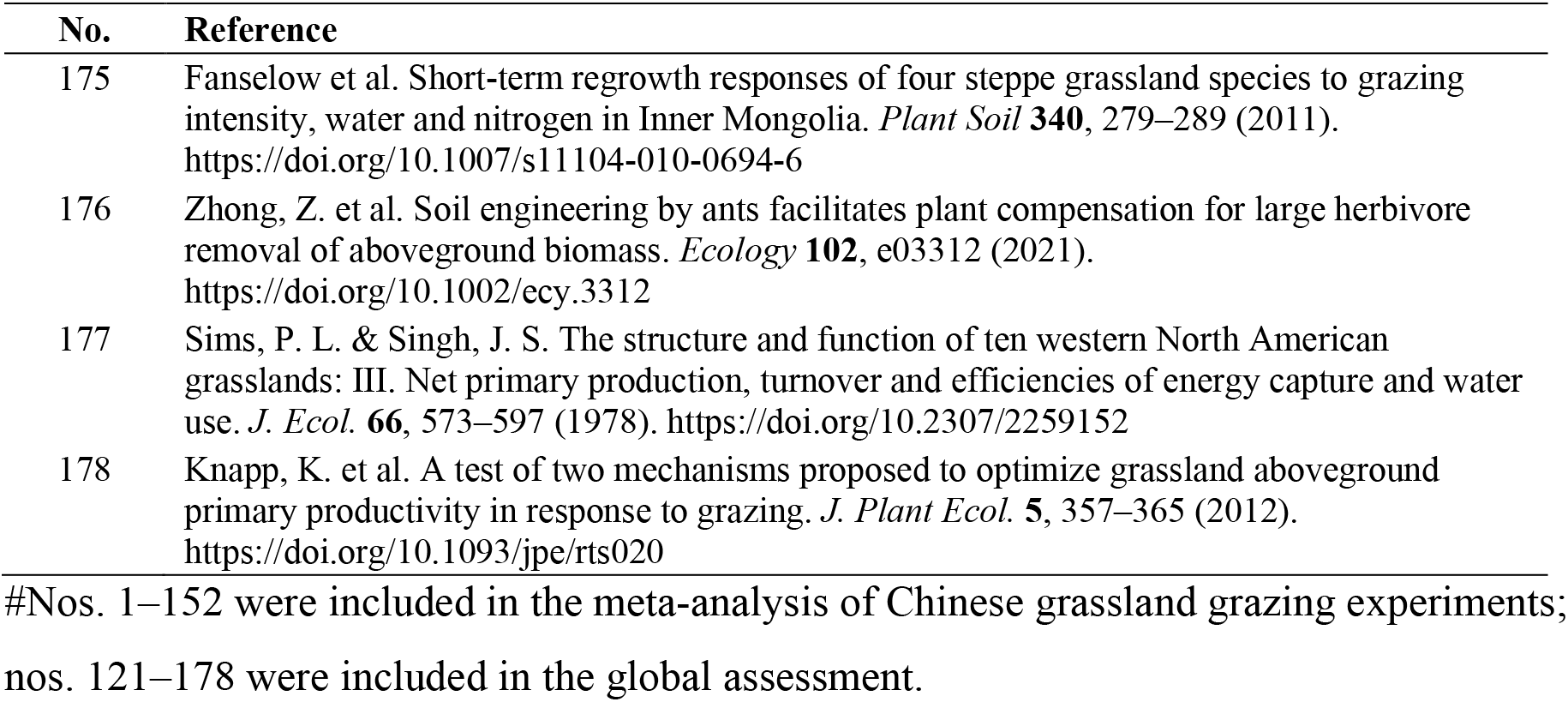
Studies included in the meta-analyses.

## References

1. Bardgett, R. D. et al. Combatting global grassland degradation. Nat. Rev. Earth Environ. 2, 720–735 (2021). 10.1038/s43017-021-00207-2

2. Briske, D. D. Rangeland systems: Processes, management and challenges (Springer Nature, 2017).

3. Strömberg, C. A. E. & Staver, A. C. The history and challenge of grassy biomes. Science 377, 592–593 (2022). 10.1126/science.add1347

4. Bai, Y. & Cotrufo, M. F. Grassland soil carbon sequestration: Current understanding, challenges, and solutions. Science 377, 603–608 (2022). 10.1126/science.abo2380

5. Ren, S. et al. Historical impacts of grazing on carbon stocks and climate mitigation opportunities. *Nat*. Clim. Change 14, 380–386 (2024). 10.1038/s41558-024-01957-9

6. Otieno, S., Njoka, J., Young, T., Mureithi, S. & Ngugi, R. Aboveground net primary productivity in grazed and ungrazed pastures: Grazing optimization hypothesis or local extinction of vegetation species. Nat. Preced. (2009). 10.1038/npre.2009.3452.1

7. Sun, Y. et al. Field-based estimation of net primary productivity and its above- and belowground partitioning in global grasslands. J. Geophys. Res. Biogeosci. 126, e2021JG006472 (2021). 10.1029/2021JG006472

8. Lauenroth, W. K. et al. Uncertainty in calculations of net primary production for grasslands. Ecosystems 9, 843–851 (2006).

9. Scurlock, J. M. O., Johnson, K. & Olson, R. J. Estimating net primary productivity from grassland biomass dynamics measurements. Glob. Change Biol. 8, 736–753 (2002). 10.1046/j.1365-2486.2002.00512.x

10. Sala, O. E. & Austin, A. T. Methods of estimating aboveground net primary productivity. in Methods in Ecosystem Science (eds Sala, O. E., Jackson, R. B., Mooney, H. A. & Howarth, R. W.) 31–43 (Springer, 2000). 10.1007/978-1-4612-1224-9_3

11. McNaughton, S. J. Compensatory plant growth as a response to herbivory. Oikos 40, 329– 336 (1983).

12. McNaughton, S. J. Serengeti migratory wildebeest: Facilitation of energy flow by grazing. Science 191, 92–94 (1976).

13. Belsky, A. J. Does herbivory benefit plants? A review of the evidence. Am. Nat. 127, 870– 892 (1986).

14. de Mazancourt, C., Loreau, M. & Abbadie, L. Grazing optimization and nutrient cycling: When do herbivores enhance plant production? Ecology 79, 2242–2252 (1998).

15. Ge, P. et al. Intensive rotational grazing has positive effects on productivity of rangeland. Agric. Ecosyst. Environ. 384, 109558 (2025). 10.1016/j.agee.2025.109558

16. Li, Y. et al. Precipitation and grazing intensity jointly shape plant compensatory growth and productivity in a semi-arid steppe ecosystem. Agric. Ecosyst. Environ. 393, 109834 (2025).

17. Oesterheld, M. & McNaughton, S. J. Effect of stress and time for recovery on the amount of compensatory growth after grazing. Oecologia 85, 305–313 (1991).

18. Field, C. B., Randerson, J. T. & Malmström, C. M. Global net primary production: Combining ecology and remote sensing. Remote Sens. Environ. 51, 74–88 (1995). 10.1016/0034-4257(94)00066-V

19. Yu, R., Evans, A. J. & Malleson, N. Quantifying grazing patterns using a new growth function based on MODIS Leaf Area Index. Remote Sens. Environ. 209, 181–194 (2018). 10.1016/j.rse.2018.02.034

20. Schmitz, O. J. et al. Animals and the zoogeochemistry of the carbon cycle. Science 362, eaar3213 (2018). 10.1126/science.aar3213

21. Yan, L., Zhou, G. & Zhang, F. Effects of different grazing intensities on grassland production in China: A meta-analysis. PLoS One 8, e81466 (2013). 10.1371/journal.pone.0081466

22. Shi, Y. et al. An improved approach to estimate stocking rate and carrying capacity based on remotely sensed phenology timings. Remote Sens. 16, 1991 (2024). 10.3390/rs16111991

23. Singh, J. S., Lauenroth, W. K. & Steinhorst, R. K. Review and assessment of various techniques for estimating net aerial primary production in grasslands from harvest data. Bot. Rev. 41, 181–232 (1975). 10.1007/BF02860829

24. Song, W. et al. Plant compensatory growth and optimal grazing intensity of grasslands in northern China: A meta-analysis of grazing experiments. Chin. Sci. Bull. 68, 1330–1342 (2023). 10.1360/TB-2022-0967

25. Hilbert, D. W., Swift, D. M., Detling, J. K. & Dyer, M. I. Relative growth rates and the grazing optimization hypothesis. Oecologia 51, 14–18 (1981). 10.1007/BF00344645

26. Zhang, Z. et al. Comprehensive analysis of grazing intensity impacts alpine grasslands across the Qinghai-Tibetan Plateau: A meta-analysis. Front. Plant Sci. 13, 1083709 (2023).

27. Ferraro, D. O. & Oesterheld, M. Effect of defoliation on grass growth. A quantitative review. Oikos 98, 125–133 (2002). 10.1034/j.1600-0706.2002.980113.x

28. Li, Y. et al. Rotational grazing promotes grassland aboveground plant biomass and its temporal stability under changing weather conditions on the Qinghai-Tibetan Plateau. Land Degrad. Dev. 31, 2662–2671 (2020). 10.1002/ldr.3596

29. Wolf, J., Chen, M. & Asrar, G. R. Global rangeland primary production and its consumption by livestock in 2000–2010. Remote Sens. 13, 3430 (2021).

30. Augustine, D. J. & McNaughton, S. J. Interactive effects of ungulate herbivores, soil fertility, and variable rainfall on ecosystem processes in a semi-arid savanna. Ecosystems 9, 1242–1256 (2006).

31. Oesterheld, M. & McNaughton, S. J. Effect of stress and time for recovery on the amount of compensatory growth after grazing. Oecologia 85, 305–313 (1991).

32. Piipponen, J. et al. Global trends in grassland carrying capacity and relative stocking density of livestock. Glob. Change Biol. 28, 3902–3919 (2022). 10.1111/gcb.16174

33. Wang, Y. et al. Grassland changes and adaptive management on the Qinghai-Tibetan Plateau. Nat. Rev. Earth Environ. 3, 668–683 (2022). 10.1038/s43017-022-00330-8

34. Chang, J. et al. Climate warming from managed grasslands cancels the cooling effect of carbon sinks in sparsely grazed and natural grasslands. Nat. Commun. 12, 118 (2021). 10.1038/s41467-020-20406-7

35. Borer, E. T. et al. Nutrients cause grassland biomass to outpace herbivory. Nat. Commun. 11, 6036 (2020). 10.1038/s41467-020-19870-y

36. Li, Y. et al. Rotational grazing promotes grassland aboveground plant biomass and its temporal stability under changing weather conditions on the Qinghai-Tibetan Plateau. Land Degrad. Dev. 31, 2662–2671 (2020).

37. Schönbach, P. et al. Grassland responses to grazing: Effects of grazing intensity and management system in an Inner Mongolian steppe ecosystem. Plant Soil 340, 103–115 (2011).

38. Nagai, S., Nasahara, K. N., Akitsu, T. K., Saitoh, T. M. & Muraoka, H. Importance of the collection of abundant ground-truth data for accurate detection of spatial and temporal variability of vegetation by satellite remote sensing. in Biogeochemical Cycles: Ecological Drivers and Environmental Impact 223–244 (2020).

39. Steier, J., Goebel, M. & Iwaszczuk, D. Is your training data really ground truth? A quality assessment of manual annotation for individual tree crown delineation. Remote Sens. 16, 2786 (2024).

40. Yuan, Q. et al. Deep learning in environmental remote sensing: Achievements and challenges. Remote Sens. Environ. 241, 111716 (2020). 10.1016/j.rse.2020.111716

41. Congalton, R. G. & Green, K. Assessing the accuracy of remotely sensed data: Principles and practices (CRC Press, 2019).

42. Steier, J., Goebel, M. & Iwaszczuk, D. Is your training data really ground truth? A quality assessment of manual annotation for individual tree crown delineation. Remote Sens. 16, 2786 (2024). 10.3390/rs16152786

43. Gao, H. et al. The carrying pressure of livestock is higher than that of large wild herbivores in Yellow River source area, China. Ecol. Model. 431, 109163 (2020). 10.1016/j.ecolmodel.2020.109163

44. Yang, F. et al. Effect of large wild herbivore populations on the forage-livestock balance in the source region of the Yellow River. Sustainability 10, 340 (2018). 10.3390/su10020340

45. Wang, H. et al. Alpine grassland plants grow earlier and faster but biomass remains unchanged over 35 years of climate change. Ecol. Lett. 23, 701–710 (2020). 10.1111/ele.13474

46. Liu, H., et al. Shifting plant species composition in response to climate change stabilizes grassland primary production. Proc. Natl Acad. Sci. USA 115, 4051–4056 (2018). 10.1073/pnas.1700299114

47. Zhan, T., Sun, J., Zhang, Z. & Liu, M. Effects of grazing on temperate grassland ecosystem based on a meta-analysis. Chinese Journal of Eco-Agriculture 28, 1847–1858 (2020). 10.13930/j.cnki.cjea.200383

48. Cao, Y., Wu, J., Zhang, X., Niu, B. & He, Y. Comparison of methods for evaluating the forage-livestock balance of alpine grasslands on the Northern Tibetan Plateau. J. Resour. Ecol. 11, 272–282 (2020). 10.5814/j.issn.1674-764X.2020.03.004

49. Nelson, J. P. Meta-analysis: Statistical methods. in Benefit Transfer of Environmental and Resource Values Vol. 14 (eds Johnston, R., Rolfe, J., Rosenberger, R. & Brouwer, R.) 329–357 (Springer, 2015). 10.1007/978-94-017-9930-0_15

50. Olson, D. M. et al. Terrestrial ecoregions of the world: A new map of life on Earth. BioScience 51, 933–938 (2001). 10.1641/0006-3568(2001)051[0933:TEOTWA]2.0.CO;2

51. Vicente-Serrano, S. M., Beguería, S. & López-Moreno, J. I. A multiscalar drought index sensitive to global warming: The standardized precipitation evapotranspiration index. J. Clim. 23, 1696–1718 (2010). 10.1175/2009JCLI2909.1

52. Spawn, S. A., Sullivan, C. C., Lark, T. J. & Gibbs, H. K. Harmonized global maps of above and belowground biomass carbon density in the year 2010. Sci. Data 7, 112 (2020). 10.1038/s41597-020-0444-4

53. Parente, L. et al. Annual 30-m maps of global grassland class and extent (2000–2022) based on spatiotemporal Machine Learning. Sci. Data 11, 1303 (2024). 10.1038/s41597-024-04139-6

